# DAMP-inducing Peptide Nanofibers and PAMP Combination Adjuvants Boost Functional Lung Tissue-resident Memory CD4^+^ T Cell Responses

**DOI:** 10.1101/2024.08.28.610131

**Authors:** Megan A. Files, Anirban Das, Darren Kim, Jeremy Buck, Janice J. Endsley, Jai S. Rudra

## Abstract

Vaccine adjuvants are typically composed of pathogen-associated molecular patterns (PAMPs) or danger-associated molecular patterns (DAMPs) that activate innate immune cells. Advances in basic immunology have demonstrated the need for various ‘types’ of protective immunity, which are difficult to achieve with a single adjuvant. The FDA approval of multiple PAMP-DAMP combinations for clinical use has led to an increased momentum in the area in recent years. Here we report the use of DAMP-inducing peptide nanofibers (PNFs) and CL429 (PAMP) combinations as subunit boosters for Bacille Calmette-Guérin (BCG). We demonstrate that pulmonary boosting with PNFs and CL429 enhances the lung-resident memory phenotype, effector cytokine profiles, and transcription factor bias of antigen-specific CD4^+^T cell populations compared to PNFs alone. Importantly, the combination significantly improved the frequency of tissue-resident memory T (T_RM_) cells which, have been shown to provide superior protection compared to circulating memory T cells. Interestingly, the T helper (Th) subset profile was driven in part driven by the route of vaccination resulting in a Th17 bias via a mucosal route or a Th1 bias when delivered intravenously. We show that following pulmonary administration, lung-resident antigen presenting cells (APCs) efficiently internalize PNFs and upregulate important co-stimulatory markers that drive T cell priming and activation. Our findings suggest that heterologous booster vaccines composed of DAMP-inducing PNFs and PAMP combinations can engage innate and adaptive immunity for generating T_RM_ cells that protect against TB and potentially other respiratory diseases.

## INTRODUCTION

Globally, tuberculosis (TB) is the leading cause of death by an infectious disease caused by *Mycobacterium tuberculosis* (*Mtb*). Despite a century of immunization with Bacille Calmette-Guérin (BCG), the only licensed *Mtb* vaccine, an estimated 10.6 million people fell ill and 1.3 million died of TB in 2022^1^. The efficacy of BCG is highly variable (0-80%) and boosting with BCG is not currently recommended^2^. Most of the world’s population is vaccinated with BCG and an attractive strategy against TB is to boost BCG-primed immunity with heterologous subunit vaccines. Most TB vaccines currently in clinical trials are subunit BCG boosters. However, a few live attenuated vaccines based on modified BCG are also in the developmental pipeline^3^.

Several emerging studies have provided valuable insight into the components of the lung immune landscape that promote protection against *Mtb*^4,5^. BCG is administered via the intradermal (i.d.) route in the clinic which generates central and effector memory populations that protect in childhood and wane by adolescence ^6,7^. However, mucosal delivery of BCG generates a subset of long-lived lung-resident CD4^+^ T cells, known as tissue-resident memory T (T_RM_) cells which, provide superior protection compared to circulating memory T cells^8^. T_RM_ cells positioned in the lung mucosa can respond to infection expediently and coordinate local immune action^9–11^. Importantly, pulmonary immunization can also generate T cell responses in distal mucosal tissues. In non-human primates, intravenous (i.v.) BCG administration afforded near complete protection from *Mtb* challenge which correlated with the accumulation of CD4^+^ T_RM_ cells in the lung^4^. While both pulmonary and i.v. delivery of BCG are known to generate superior immunity, neither can be practically nor safely employed in the human population. Thus, the development of mucosal vaccines that boost immunity and augment lung CD4^+^ T_RM_ cells following a BCG prime holds great promise for protection against TB^12–14^.

Current vaccines in the TB clinical development pipeline can be broadly divided into adjuvanted protein subunit vaccines or viral-vectored vaccines. Recently, immunoengineering approaches based on polymeric micro/nano particles, liposomes, and virus-like particles (VLPs) have also shown promise in preclinical models of *Mtb*^15^. Our lab studies self-assembling peptide nanofibers (PNFs) as adjuvant-free vaccine delivery vehicles and we demonstrated their utility in multiple preclinical models of disease^16^. Transcriptomic analysis of nanofiber treated dendritic cells (DCs) suggested that the adjuvanticity is in part due to the release of damage-associated molecular patterns (DAMPs). The PNF platform, due to its synthetic and modular nature, offers the customization of immune responses through the stoichiometric inclusion of antigens and adjuvants^17^. This is highly promising for the installation of CD4^+^ T_RM_ cells in the lungs and other mucosal tissues for protection against respiratory pathogens like *Mtb*. In this study, we sought to determine the effects of combining PAMP (pathogen associated molecular pattern) agonists with DAMP-inducing PNFs on the functional profile of lung T_RM_ cells.

To address this, we generated fusion peptides composed of the self-assembling peptide KFE8 (FKFEFKFE) and Ag85B (FQDAYNAAGGHNAVF), a protective CD4^+^ T cell epitope from Ag85B and a component of multiple TB vaccines in clinical trials^18^. We boosted BCG-primed mice with KFE8-Ag85B nanofibers with or without CL429, a dual agonist of toll-like receptor 2 (TLR2) and nucleotide-binding oligomerization domain-containing protein 2 (NOD2) via the pulmonary or i.v. route. The adjuvanting effects of PNFs has not been tested via the i.v. route to date. We also determined the characteristics of lung antigen presenting cells (APCs) responsible for priming adaptive immunity. Our data shows that KFE8-Ag85B nanofibers are taken up by a myriad of lung APCs and promote surface expression of co-stimulatory molecules. To assess the effects of PNF-CL429 combination adjuvants, we probed memory precursor effector cells (MPEC) for their expression of transcription factors EOMES, GATA3, RORγt, and T-bet that regulate Th differentiation. We found superior expansion of antigen-specific effector T cell populations with potential to become memory T cells. Interestingly, we found route-specific differences with a pulmonary boost directing a Th17 bias whereas an i.v. boost led to a Th1 bias. These results were supported by cytokine profiles in lung supernatants following antigen recall. Using Isoplexis technology^19^, we assessed the single cell secretome profiles of lung memory CD4^+^ T cells for polyfunctional cytokine production.

Collectively, our results suggest a promising vaccine platform for generating BCG boosters that generate lung CD4^+^ T_RM_ cells that could potentially protect against TB. These findings suggest that DAMP-PAMP adjuvant combinations can augment T_RM_ responses and that different routes of boosting could direct the expansion of memory T cell populations towards different phenotypes. These results set the stage to advance the development of multivalent, nanofiber subunit nano vaccines against *Mtb* that would expand T_RM_ cells as part of a heterologous prime-boost strategy in combination with BCG.

## MATERIALS AND METHODS

### Peptide synthesis and nanofiber preparation

Peptides KFE8 (Ac-FKFEFKFE-Am), KFE8-Ag85B (FKFEFKFE-GGAAY-FQDAYNAAGGHNAVF) and fluorescently labeled Cy5-KFE8-Ag85B were purchased from GenScript (NJ, USA). GGAAY is a cleavable linker which acts as a spacer between KFE8 and Ag85B. Fibrils of KFE8-Ag85B and Cy5-KFE8-Ag85B were prepared by resuspending the lyophilized peptide in sterile, endotoxin free water to a stock concentration of 1 mM solution. The stocks were further diluted to 0.1 mM in sterile PBS just prior to use.

### Transmission Electron Microscopy

KFE8 or KFE8-Ag85B solutions (0.1 mM in water) were applied to 200 mesh, carbon-coated, copper grid (10 μL) for 2 minutes. Excess sample was blotted and the grids were stained with uranyl formate for 1 min and rinsed thrice in water. Bright field images were taken with a JEOL JEM-2100 scanning transmission electron microscope (STEM) with an accelerating voltage of 120 kV at Washington University Center for Cellular Imaging.

### Circular Dichroism (CD) Spectroscopy

CD spectra of the peptide solutions (0.1 mM) were recorded on a Jasco J-815 circular dichroism spectrometer. Spectra of the peptide solutions (3 scans) were collected at 20°C in a 1 mm quartz cuvette between 260 and 190 nm with a bandwidth of 1 nm, 0.5 nm step, and 2 sec averaging time per step. Solvent background was subtracted.

### Thioflavin-T (ThT) and ANS (8-anilo-1-napthalenesulfonic acid) assays

Peptide fibrils (50 µM in water) were mixed with ThT (50 µM in water) and fluorescence emission was measured (470-700 nm) upon excitation (440 nm) using a Synergy H1 microplate reader (BioTek). For ANS assays, peptide solutions (20 µM) were mixed with ANS (250 µM) and emission spectra was collected (380-700 nm) following excitation at 350 nm. Intrinsic peptide fluorescence was measured and subtracted as background.

### Bacteria and growth conditions

BCG Pasteur (ATCC 35734) was a kind gift from Dr. Christina Stallings at Washington University in St. Louis. The culture was propagated using Middlebrook 7H9 media supplemented with 10% v/v OADC (oleic acid, albumin, dextrose, catalase), 0.05% Tween80 v/v, and 0.5% v/v glycerol and grown a 37°C incubator. For vaccination, cultures were grown to an optical density (OD600) of 0.4-0.6, pelleted, and washed twice in PBS. Bacterial pellets were resuspended in sterile PBS at a final density of 5.5 x 10^6^ colony-forming units per 1 mL (CFU/mL). Experiments with BCG were conducted under BSL2 or ABSL2 conditions with appropriate safety equipment and following approved institutional protocols.

### Animals and immunizations

C57BL/6 mice (6-8 weeks old females) were acquired from Jackson Labs and housed in an ABSL2 facility. All studies were approved by the Institutional Animal Care and Use Committees at the University of Texas Medical Branch or at Washington University in St. Louis. Mice were given ad libitum access to food and water. For BCG inoculations, mice were given intradermal (i.d.) vaccinations of 5.5 x 10^5^ CFU BCG in 100 µL of sterile PBS. Nanofiber boosts were given via the i.t. or i.v. route under 1-3% isoflurane anesthesia in 100 µL of sterile PBS containing 100 µM KFE8-Ag85B with or without 10 ng of CL429 (n = 4-5/group or n = 9/group). For uptake studies by lung APCs, 100 µL of sterile PBS containing 100 µM Cy5-KFE8-Ag85B was administered via the i.t. route under anesthesia and mice were euthanized at 4h or 24h and lungs were extracted for further processing. Control mice received PBS only.

### Tissue processing

In the experiments described in Figures 1-8, lung tissue was minced and incubated in 3 mL of serum-free media containing 0.5 mg/mL DNase I (Roche, 10104159001) and 1 mg/mL collagenase type IV (Worthington, LS004188) in a 37°C humidified incubator for ∼1h. Spleen tissue was dissociated using a MACS system (Miltenyi Biotec, USA). The digests were passed through a 70 µm cell strainer into 10 mL of complete RPMI media supplemented with 10% FBS and 1% penicillin-streptomycin (Pen-Strep). Cells were pelleted at 300 × g for 5 minutes and RBC lysis buffer (Sigma, R7757-100ML) was added for one minute followed by quenching with PBS. Cells were washed once and half of the lung cells were stained for flow cytometric analysis. The remaining half were used for single cell secretome analysis. Similar to lung lymphocytes, CD4^+^ T cells were isolated from half each disrupted spleen, and the remaining half was frozen in 90% FBS and 10% DMSO and stored in liquid nitrogen until further use.

**Figure 1.**
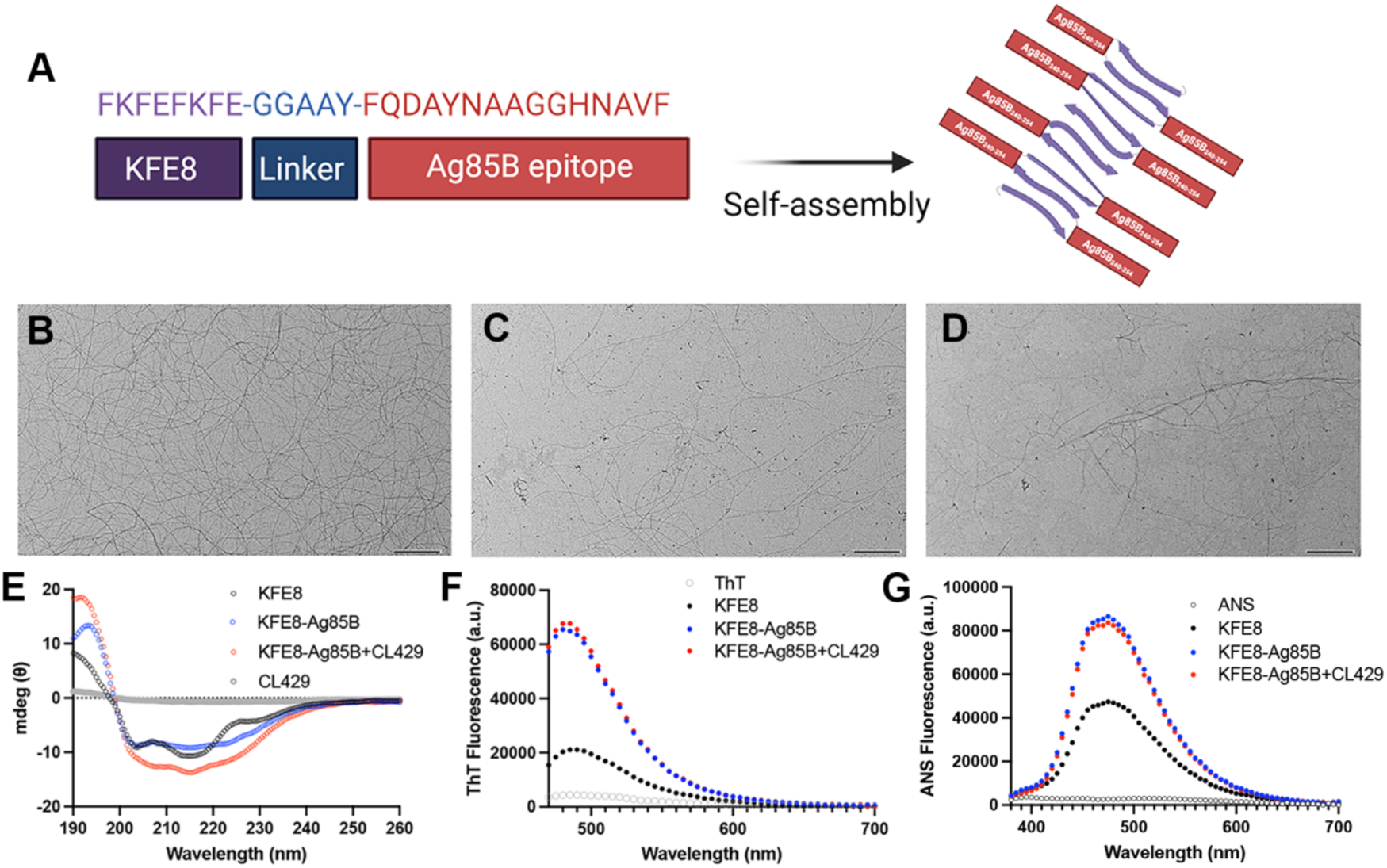
Ag85B-KFE8 construct self-assemble into β-sheet rich nanofibers. Schematic showing the fusion of Ag85B epitope to the self-assembling domain KFE8 via GGAAY linker (**A**). Representative TEM images of KFE8 (**B**), Ag85B-KFE8 (**C**) and Ag85B-KFE8+CL429 (**D**) (0.1 mM in water, scale bar is 500 nm). Secondary structure analysis and circular dichroism data showing a β-sheet dominant spectrum as evidenced by the minima at 215 nm (**E**). ThT fluorescence spectra showing enhanced β-sheet content in Ag85B-KFE8 compared to KFE8 (**F**). ANS fluorescence spectra showing increased hydrophobicity in Ag85B-KFE8 (G). Schematic was created in BioRender.com.

### Cell staining protocols

Lung cell suspensions prepared above were resuspended in 1 mL of PBS containing Zombie Green Viability Dye (0.1% Biolegend, 423112) and incubated at 4°C for 15 min. Cells were washed in PBS, resuspended, and incubated with Fc block (BD Bioscience, 553142) in cell staining buffer (1% FBS v/v, 0.1% sodium azide in Ca^2+^ and Mg^+2^ free PBS) for 5 minutes. To identify Ag85B-specific CD4^+^ T cells and characterize the cellular phenotype of lung lymphocytes, cells were stained for 30 min at room temperature with an MHC-II Ag85B tetramer conjugated to streptavidin-PE (MBL, TS-M719-1) followed by an additional hour of staining at 4°C with anti-CD3 (AlexaFluor 350, R&D Biosystems, FAB484U-100UG), anti-CD4 (BUV496, BD Biosciences, 564667), anti-CD8α (AlexaFluor 700, Biolegend, 100730), anti-CD44 (PerCP-Cy5.5, Biolegend, 103032), anti-CD62L (PE-Dazzle 594, Biolegend, 104448), anti-CCR7 (Brilliant Violet 605, Biolegend, 120125), anti-CD69 (Brilliant Violet 421, Biolegend, 104528), anti-KLRG1 (BUV 737, BD Biosciences, 741812), anti-CXCR3 (Brilliant Violet 480, BD Biosciences, 746651), anti-CD127 (PE-Vio770, Miltenyi Biotec, 130-117-782), anti-CX3CR1 (APC-Fire 750, BioLegend, 149040), and anti-PD-1 (SuperBright 645, Thermo-Fisher Scientific, 64-9985-82) antibodies. Using the eBioscience Foxp3/Transcription Factor Staining Buffer Set (Invitrogen, 00-5523-00), cells were then permeabilized in 1 mL fixation/permeabilization solution and incubated for 30 minutes at 4°C. Cells were washed two times in permeabilization buffer and centrifuged 500 ×g for 5 minutes, and resuspended in an antibody cocktail containing anti-RORγt (Alexa Fluor 647, BD Biosciences, 562682), anti-T-bet (Brilliant Violet 785, Biolegend, 644835), anti-GATA3 (Brilliant Violet 711, BD Biosciences, 565449), and anti-EOMES (PE-Cy5, Thermo Fisher Scientific, 15-4875-82) antibodies to stain at room temperature for 30 minutes. Cells were then washed in permeabilization buffer and fixed in 2% ultrapure formaldehyde (Fisher Scientific, 50-980-487). Acquisition of samples was performed using BD LSRFortessa X-20 flow cytometer at Washington University in St. Louis in the Bursky Center for Human Immunology & Immunotherapy Programs. The compensation matrix was developed using UltraComp ebeads (Invitrogen, 01-2222-42) and calculated using the BD FACS Diva software.

For experiments detailed in Figure 9, mice were given i.t. immunizations of Cy5-KFE8-Ag85B and lungs were collected at 4 and 24 hours after administration. Lung tissue was minced and digested as described above. After RBC lysis, washing, and centrifugation, cells were resuspended in PBS containing 0.1% v/v Zombie Green Viability Dye and incubated at 4°C for 15 minutes. Following a wash in PBS and incubation in Fc Block, cells were incubated at 4°C for 1h in an antibody cocktail containing anti-CD11b (PE, Biolegend, 101207), anti-CD11c (Brilliant Violet 421, Biolegend, 117330), anti-CD206 (Brilliant Violet 650, Biolegend, 141723), anti-F4/80 (PerCP-Vio700, Miltenyi Biotec, 130-118-466), anti-CD103 (Brilliant Violet 605, Biolegend, 121433), anti-CD80 (APC-Vio770, Miltenyi Biotec, 130-116-399), anti-CD86 (Brilliant Violet 785, Biolegend, 105043), and anti-CD73 (PE-Dazzle 594, Biolegend, 127233) antibodies. Cells were resuspended and in 2% ultra-pure formaldehyde prior to acquisition on an ACEA NovoCyte 3000 and the compensation matrix was made using Ultra Comp beads and the FlowJo (v10) software. Tissue preparation for data shown in Figures 10-13 were performed as described for Figure 9. Viable cells were identified using live/dead fixable near IR stain kit (Invitrogen, L34976). To characterize APC populations associating with KFE8, cells were stained with an antibody cocktail containing anti-SiglecF (BUV395, BD Biosciences, 740280), anti-CD11b (SuperBright600, Thermo-Fisher, 63-0112-82), anti-CD11c (PE-Dazzle 594, Biolegend, 117348), anti-Ly6G (Pacific Blue, Biolegend, 127612), anti-CD64 (FITC, Biolegend, 139315), anti-MHCII (Alexa Fluor 700, eBiosciences, 56-5321-82), and anti-F4/80 (Brilliant Violet 510, Biolegend, 123135) antibodies at 4°C for 1 hour. Samples were acquired using a BD Fortessa LSRII at the UTMB Flow Cytometry and Cell Sorting Facility. The compensation matrix was calculated using the Diva FACS software.

### Single-cell secretome and supernatant analysis

Lungs or spleens collected from experiments in figures 1-8 were purified to isolate CD4^+^ T lymphocytes using the EasySep Mouse CD4^+^ T Cell Isolation Kit (STEMCELL, 19852) and the EasySep Magnet (STEMCELL, 18102) according to manufacturer’s instructions. To ensure sufficient CD4^+^ T cell numbers, lungs from three mice in each group were pooled, resulting in three samples representing each treatment group. CD4^+^ T cells isolated from lung were stimulated with anti-CD3e (Thermo-Fisher, 16-0031-86) and anti-CD28 (Thermo-Fisher, 16-0281-85) antibodies in complete RPMI supplemented with 55 μM 2-mercaptoethanol (2-ME) for 48 hours in a humidified incubator at 37°C and 5% CO_2_. For cytokine and secretome analysis, stimulated cells were collected from plates and pelleted at 300 × g for 10 minutes. The supernatants were stored at -80°C for analysis. A cell membrane stain provided by the manufacturer was prepared and added (100 μL/10^6^ cells) to each sample tube. After incubating for 10 minutes at 37°C in the dark, cells were stabilized with 5 volumes of complete RPMI, washed, and resuspended to a final cell density of 10^6^ cells/mL. IsoCode chips (Single-Cell Adaptive Immune Chip (M), ISOCODE-1004) were loaded with 30 μL of cell suspension and inserted into the IsoLight instrument. Supernatants were analyzed in duplicate for IFN-γ, IL-2, IL-17A, TNF-α, IL-4, IL-5, IL-13, IL-6, and IL-1β using multiplex ELISA (R&D Systems, LXSAMSM-09). Supernatants are representative of 3 mice per sample. Two samples per treatment group were analyzed for cytokines.

### Antigen recall assay

Bone marrow-derived dendritic cells were generated as previously described. Briefly, bone marrow was harvested from C57/B6 mice, treated with RBC lysis buffer, and plated incubated in complete media containing 10 ng/mL mouse rGM-CSF and 5 ng/mL mouse rIL-4 for 7 days. BMDCs were then plated at a concentration of 1 × 10^6^ cells per well in a 96-well plate and incubated with either Ag85B or PBS. Thawed splenocytes were added to the 96-well plate at a concentration of 1×10^7^ cells/well and incubated for 72h. At 6h prior to harvesting cells, Golgistop was added according to manufacturer’s instructions. The 96-well plates were centrifuged at 300 x g and supernatants were collected and stored at -80°C. Cells were washed and stained for viability, then stained with fluorescent antibodies targeting surface molecules CD3, CD4, and CD8. After surface staining, cells were permeabilized using BD Fix Perm and cells were stained intracellularly with antibodies targeting mouse IFN-γ, TNF, IL-2, and IL-17A. After washing in PBS and fixing cells in 2% paraformaldehyde, cells were acquired using an LSR Fortessa and flow cytometric data was analyzed using FlowJo v10.

### Statistical analysis

Data were analyzed using GraphPad Prism version 9 and presented as mean (± SEM). Significant differences were determined using a one-way ANOVA followed by an appropriate ad hoc test for differences due to treatment, or between treatment groups, as indicated in each figure legend. Statistical analysis of data sets containing only two experimental groups was conducted using a two-tailed unpaired t test. Data shown in Figure S6 was plotted and extracted from the IsoSpeak software 2.9.0. p-values of <0.05 were considered significant.

## RESULTS

### KFE8-Ag85B fusion peptide self-assembles into β-sheet rich nanofibers

To generate Ag85B nanofibers, the immunodominant *Mtb* epitope Ag85B (FQDAYNAAGGHNAVF) was coupled to the self-assembling peptide KFE8 (FKFEFKFE) via a cleavable linker (GGAAY) (**Fig. 1A**). In physiological buffers, the fusion peptide (KFE8-GGAAY-Ag85B) assembled into nanofibers with KFE8 peptide forming the core and Ag85B arrays on the surface **(Fig. 1A**). TEM micrographs showed nanofibers (7-8 nm) for KFE8 (**Fig. 1B**) and KFE8-Ag85B (**Fig. 1C**) indicating self-assembly and presence of CL429 did not disrupt fibril morphology (**Fig. 1D**). CD spectra verified that the secondary structure was preserved as evidenced by the double absorbance minima at 205 and 220 nm (**Fig. 1E**). These data are consistent with prior studies where KFE8 was amenable to functionalization with peptide epitopes while maintaining morphology and secondary structure. Further, appending the Ag85B epitope to KFE8 enhanced β-sheet fibril formation with an increase in ThT fluorescence for KFE8-Ag85B compared to KFE8 (**Fig. 1F**). The net hydrophobicity of the constructs was assessed using ANS which binds to exposed hydrophobic patches or cavities and data indicated enhanced fluorescence emission for KFE8-Ag85B compared to KFE8 (**Fig. 1G**). Together, these data show that conjugation of Ag85B epitope to KFE8 results in the robust formation of β-sheet rich nanofibers and there was no significant impact on fibril formation or secondary structure in the presence of CL429.

### Boosting with KFE8-Ag85B PNFs and CL429 augments effector T cell populations

Mice were primed with BCG and boosted twice via the i.t. or i.v. route with KFE8-Ag85B PNFs with or without CL429 (**Fig. 2A**). Mice were humanely euthanized one week after the second boost and lungs and spleen were collected to assess effector and memory T cell subsets via multiparametric flow cytometry. Cells from lung homogenate were gated on CD4^+^ CD3^+^ lymphocytes or CD8^+^ CD3^+^ lymphocytes. Central and effector memory cells were characterized as CD44^hi^ CD62L^hi^ and CD44^hi^ CD62L^lo^ respectively. The T_EM_/T_EF_ subset was further evaluated for expression of CD127 (IL-7Ra) to differentiate T_EM_ (CD127^+^) from T_EF_ (CD127^-^). Gating strategy is shown in Figure S1.

**Figure 2.**
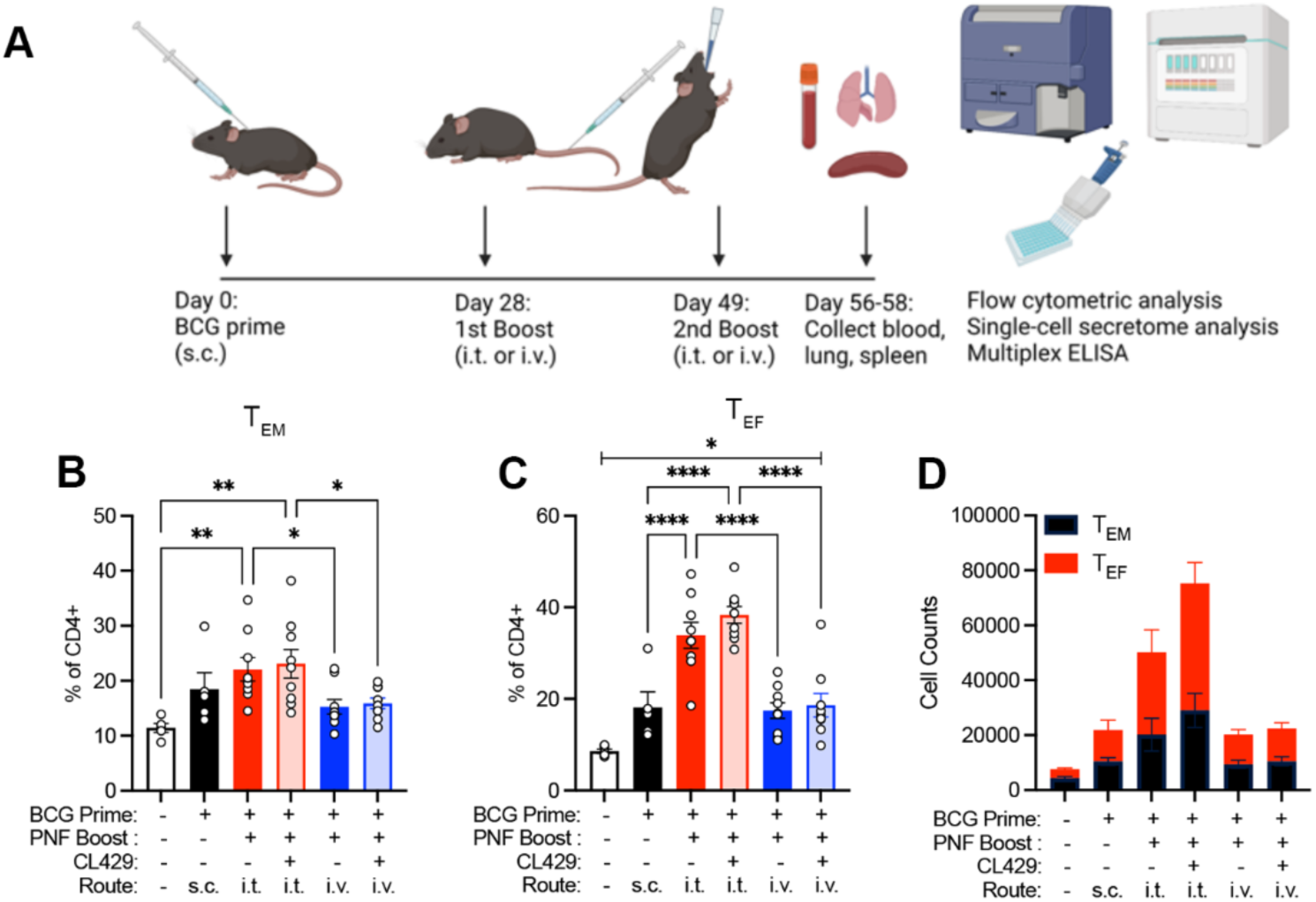
Pulmonary boost with KFE8-Ag85B in BCG-primed mice increases the frequency of CD4^+^ T_EM_ and T_EF_ cells. BCG-primed mice were given two boosts four weeks and seven weeks later with Ag85B-KFE8 nanofibers with or without CL429 adjuvant. Booster doses were delivered via the i.t. or i.v. route. Mice were euthanized after the second boost and lungs, spleen, and blood were collected for flow cytometry, single cell secretome analysis, and antigen recall assays **(A)**. CD4^+^ T_EM_ and T_EF_ cells were identified as CD44^hi^ CD62L^lo^ and further differentiated based on their expression of CD127. T_EM_ (CD127^+^) cells (**B**) and T_EF_ (CD127^-^) (**C**) cells are shown as a percentage of the total CD4^+^ T cell population. Total counts of T_EM_ and T_EF_ cells are shown in stacked plots (**D**). Statistical significance was determined by one-way ANOVA followed by a Benjamini Krieger Yuketieli test to control for multiple comparisons (n = 5 in untreated and BCG only groups, n = 9 in boosted groups). *p<0.05, **p<0.005, ****p<0.0005. Schematic was created in BioRender.com.

Quantification of flow data indicated that BCG-primed mice receiving i.t. boosts had significantly expanded populations of CD4^+^ T_EM_ and T_EF_ cells. Increased levels of T_EM_ cells (22%) compared to untreated controls (11%) were detected however, this was not different from mice primed with BCG alone (18%) (**Fig. 2B**). Inclusion of CL429 or i.v. boosting did not enhance T_EM_ cells significantly compared to untreated or BCG-primed groups (**Fig. 2B**). A significant finding in our data was the expansion of T_EF_ cells in mice receiving Ag85B PNFs via the i.t. route compared to BCG alone (34% vs 17%) (**Fig. 2C**). Importantly, this increase was route-dependent with greater expansion observed via the pulmonary route compared to the i.v. route (**Fig. 2C**). Addition of CL429 did not influence the frequency of T_EM_ or T_EF_ cells in all boosted mice. Total cell counts are shown in **Fig. 2D**, with higher numbers of CD4^+^ T_EM_ and T_EF_ cells in mice receiving i.t. boost compared to an i.v. boost.

### Boosting with nanofiber vaccines generates T_EM_ and T_EF_ cells associated with greater proliferative potential

Numerous studies have described phenotypic markers of short-lived effector cells and long-lived memory T cells^8,9,20,21^. KLRG1 (killer cell lectin-like receptor G1) has been shown to be a marker of terminally differentiated effector CD8^+^ T cells in animal models of viral infection^22,23^ as well as *Mtb* infection^24^. Although there is emerging evidence that CD4^+^T cells do not exhibit the same patterns of differentiation observed in CD8^+^T cells^25^, in the context of persistent infections, KLRG1 expression is associated with terminal differentiation. Parenchymal lung CD4^+^ T cells in the setting of *Mtb* infection are predominantly KLRG1^-^ CX3CR1^-^ and produce less IFN-γ than circulating memory populations^8^, but are highly proliferative and survive longer than their intravascular KLRG1^+^ counterparts^24^. Expression of KLRG1 on CD4^+^ T cells in humans with *Mtb* infection was shown to be increased due to antigen-driven T cell activation^26,27^, which was further supported by a report that anti-tubercular therapy reduced the expression KLRG1 in a mouse model of *Mtb* infection^28^. We therefore identified terminally differentiated T cells by the expression of KLRG1 and CX3CR1 (**Fig 3A**).

**Figure 3.**
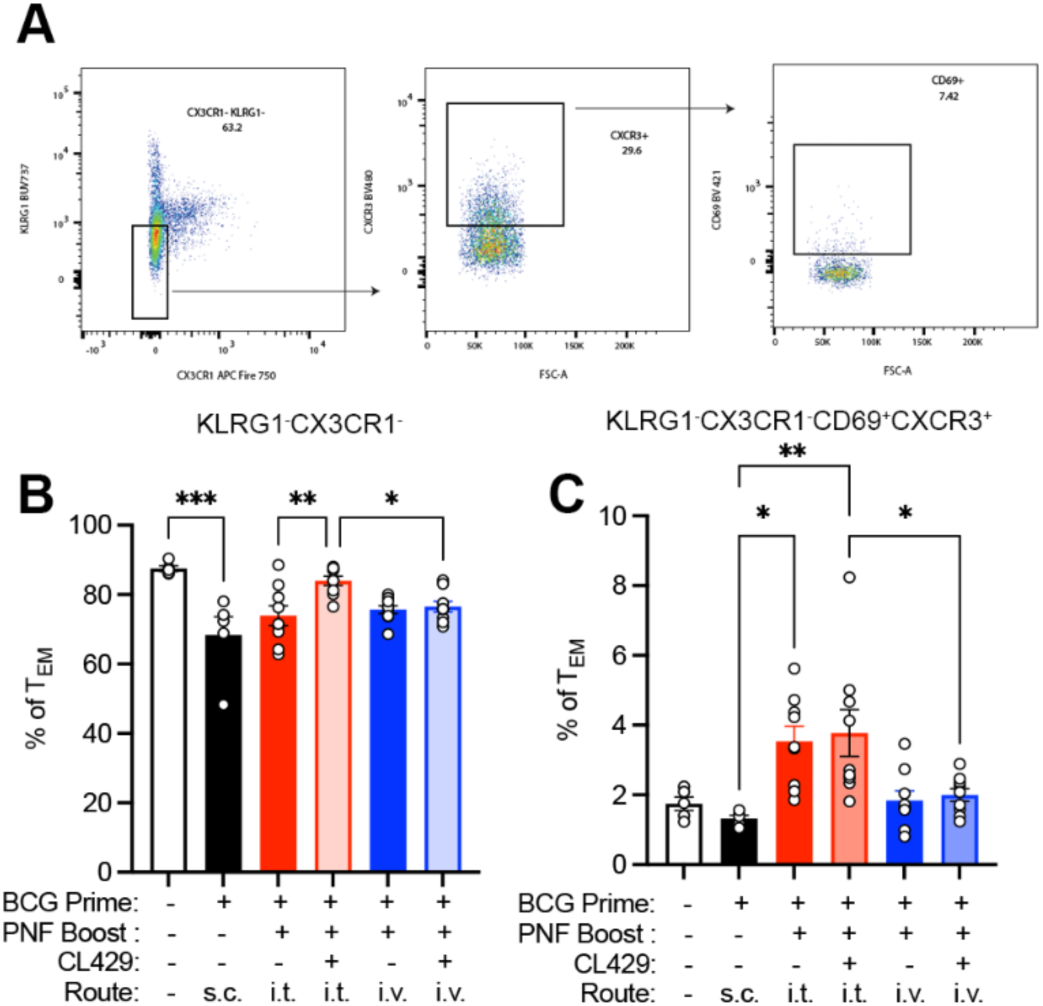
Boosting with nanofiber vaccines generates T_EM_ cells with greater proliferative potential. CD4^+^ T_EM_ T cells were analyzed for their expression of KLRG1, CX3CR1, CXCR3, and CD69 (**A**). Cells lacking expression of KLRG1 and CX3CR1 are represented as a percentage of total CD4^+^ T_EM_ population (**B**). Cells that were CXCR3^+^ CD69^+^ are represented as a percentage of KLRG1^-^CX3CR1^-^ T_EM_ cells (**C**). Significant differences were determined by one-way ANOVA followed by a Benjamini Kreiger Yuketieli test to control for multiple comparisons (n = 5 in untreated and BCG only groups, n = 9 in boosted groups). Significance was determined by one-way Brown Forsythe and Welch ANOVA to account for differences in standard deviation followed by a Benjamini Kreiger Yuketieli test to control for comparisons (n = 5 in untreated and BCG only groups, n = 9 in boosted groups). *p<0.05, **p<0.005, ***p<0.0005.

Data shows that in mice primed with BCG alone, the pool of KLRG1^-^ CX3CR1^-^ CD4^+^ T_EM_ cells is significantly reduced compared to untreated controls (**Fig. 3B**). This is presumably due to an ongoing immune response to persistent bacilli and presence of antigen. Pulmonary boosting with combined nanofiber and CL429 formulations increased the frequency of KLRG1^-^CX3CR1^-^ CD4^+^ T_EM_ in comparison to BCG alone (**Fig. 3B**). However, i.t boosting with Ag85B nanofibers was sufficient to induce a significant increase in KLRG1^-^ CX3CR1^-^ CD4^+^ T_EF_ cells which was further enhanced by addition of CL429 (**Fig. S2**). Increased expression of CXCR3, an important homing marker, and CD69, a membrane-bound type II C-lectin receptor, have been identified as important T_RM_ markers^29^. After selection of CD44^hi^ CD62L^lo^ CD127^+^ cells in the live CD3^+^ CD4^+^ gate, we identified a core population of T_RM_ cells as CX3CR1^-^KLRG1^-^CD69^+^CXCR3^+^ cells. Data indicated that boosting via the i.t. route elicited significantly higher frequencies of CD4^+^T_RM_ cells compared to mice receiving BCG only (**Fig. 3C**). Interestingly, addition of CL429 did not lead to a significant change in this population.

### Boosting BCG-primed mice with CD4^+^ T cell targeting nanofiber vaccines enhances activation of CD8^+^ T cells

Despite Ag85B being a well-characterized *Mtb* CD4^+^ T cell epitope, we also examined activation of T_EM_ and T_EF_ CD8^+^ T cell populations after boosting. Internal epitopes of FQD**AYNAA**G**GHNAVF** (bolded) were previously shown to be presented by H2^K^ MHC I of C57BL/6 mice vaccinated with *M. ulcerans*^30^. Data showed an overall increase in T_EM_/T_EF_ CD8^+^ T cell populations in i.t. boosted mice compared to the i.v. route (**Fig. S3A**). Interestingly, the T_EM_ frequency in i.t. boosted mice significantly decreased (**Fig. S3B**) while the T_EF_ frequency increased (**Fig. S3C**) but no significance was detected compared to BCG alone. Counts of T_EM_ and especially T_EF_ cells increased in all vaccinated groups, however the number of CD8^+^ T_EF_ cells was dramatically increased in mice that received an i.t. boost without CL429 (**Fig. S3D**). This is consistent with prior work from our lab where induction of antigen specific CD4^+^ T cells through boosting with nanofibers also improved CD8^+^ T cell responses^31^. Other studies have corroborated this effect as evidenced by the enhanced CD8^+^ CTL response to Ag85B^32^.

CL429 did not influence the frequency of T_EM_ or T_EF_ CD8^+^ T cells when boosted via either route. Mice boosted i.t. with Ag85B nanofibers and CL429 had a higher frequency of CX3CR1^-^ KLRG1^-^ CD8^+^ T_EM_ cells in the lungs compared with BCG prime alone (**Fig S3E**). All i.t. boosted mice demonstrated significant increase in CD8^+^ CX3CR1^-^ KLRG1^-^ T_EF_ cells and addition of CL429 significantly enhanced this population (**Fig. S3F**). Together, these data demonstrate that i.t. boosting with Ag85B nanofibers generates robust populations of effector and memory CD8^+^ T cells in the lung. Comparative analysis of the boosting routes suggests that pulmonary delivery of Ag85B nanofibers effectively targets the lung mucosal immune system and enhances memory-precursor CD8^+^ T cell populations in the lungs of mice previously immunized with BCG. Addition of CL429 in mice boosted via i.t. route also increased the frequency of T_RM_ CD8^+^ T cells. These results are exciting as they are a strong predictor of memory cell formation.

### Incorporation of CL429 adjuvant expands the pool of Ag85B-specific CD4^+^ T cells

We next evaluated the antigen specificity of CD4^+^ T populations using Ag85B tetramer staining (**Fig. 4A**). Consistent with our previous studies^33^, i.t. boosting with KFE8-Ag85B expanded the pool of tetramer-positive CD4^+^ T cells in the lungs of BCG-primed mice (**Fig 4B**). Inclusion of CL429 significantly enhanced the frequency of Ag85B-specific CD4^+^ T cells in the lungs compared with BCG when delivered via i.t. route. Further, significantly higher frequencies of CX3CR1^-^KLRG1^-^ CD69^+^CXCR3^+^ antigen-specific T_RM_ cells were detected compared to mice receiving BCG only. A small increase in antigen-specific T_RM_ cells was observed in i.v. boosted groups compared to mice receiving BCG alone (**Fig. 4C**). Tetramer-positive cells were additionally categorized as either T_CM_, T_EM_, or T_EF_ and boosting with PNF-CL429 combination adjuvants via the i.t. route instilled the highest numbers of antigen-specific CD4^+^ T cells in all three compartments compared to BCG alone (**Fig. 4D-F**). Total cell counts are shown in **Fig. 4G**. Overall, inclusion of CL429 enhances antigen-specific cell populations compared to boosting with Ag85B PNFs alone and with expression patterns associated with memory populations of memory CD4^+^ T cells.

**Figure 4.**
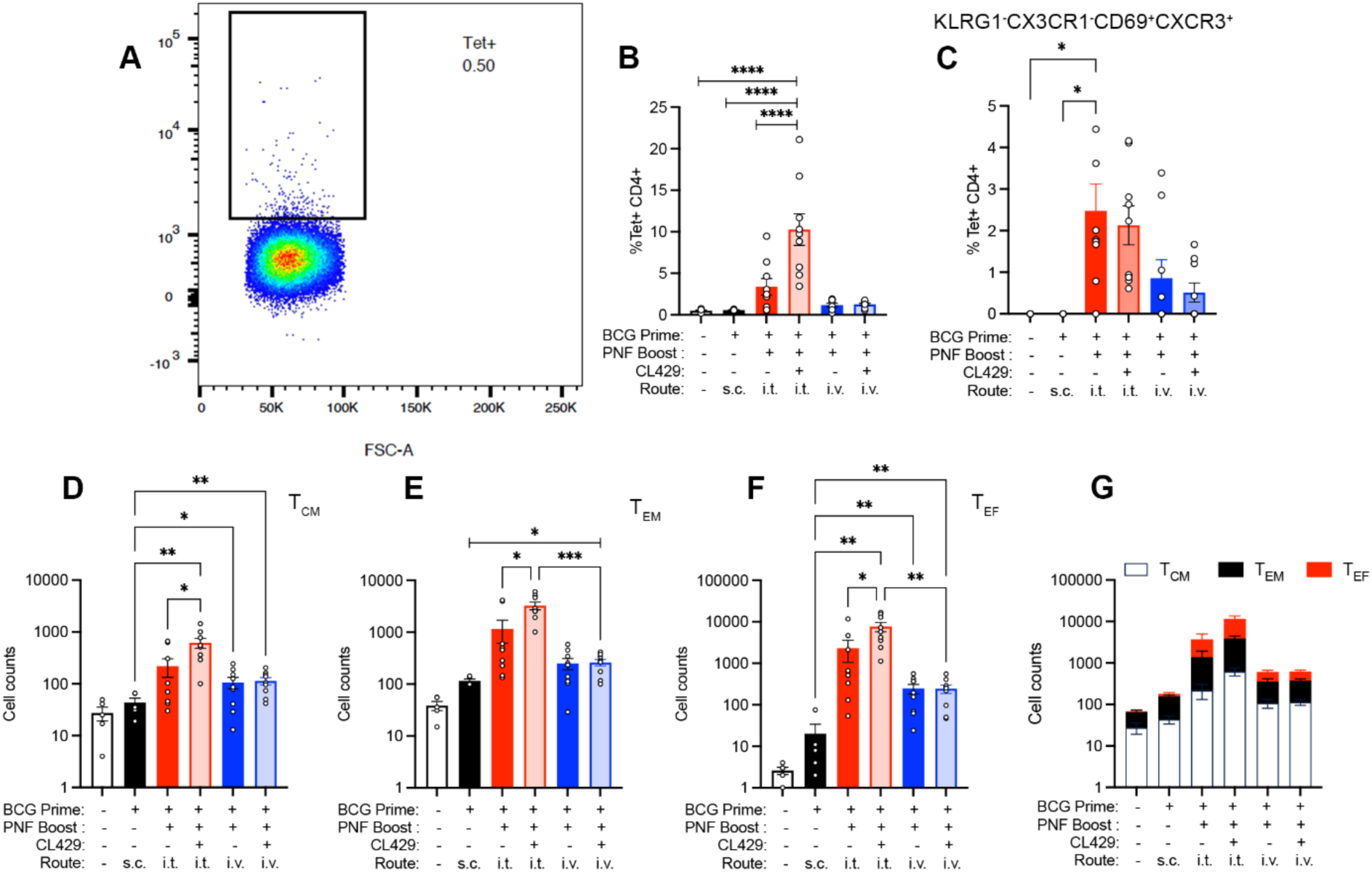
new. Addition of CL429 expands pools of Ag85B-specific CD4^+^ T cells in the lung which express markers of tissue residency. Ag85B-specific CD4+ T cells were identified following staining with PE-labeled Ag85B tetramer **(A)**. Ag85B-specific CD4^+^ T cells in the lungs of vaccinated mice are represented as a percentage of all CD4^+^ T cells (**B**). Tetramer-positive cells were analyzed for their expression of surface markers associated with long-lived T_RM_-like cells (KLRG1^-^CX3CR1^-^CXCR3^+^CD69^+^) (**C**). The total counts of tetramer-positive CD4^+^ T_CM,_ T_EM,_ and T_EF_ are shown individually **(D-F)** and as a stacked plot **(G)**. Significance was determined by one-way Brown Forsythe and Welch ANOVA to account for differences in standard deviation followed by a Benjamini Kreiger Yuketieli test to control for comparisons (n = 5 in untreated and BCG only groups, n = 9 in boosted groups). *p<0.05, **p<0.005, ***p<0.001.

### Boosting with combination adjuvants drives transcription factor bias in CD4^+^ T cells in a route-dependent fashion

We next determined the Th subsets that are promoted following i.t. or i.v. boost with combination adjuvants. The expression of transcription factors EOMES, GATA3, RORγT, and T-bet were analyzed in tetramer-positive CD4^+^ T cells and data indicated a route- and adjuvant-dependent effect (**Fig. 5**). EOMES is a T-box transcription factor which is important for the function and homeostasis of effector and memory T cells. However, overexpression of EOMES can lead to T cell exhaustion and is associated with a bias toward a Th1 phenotype and suppression of Th17^34^. A comparative reduction of EOMES was observed in all boosted groups, most significantly in mice boosted with the combination adjuvant via the i.t. route (**Fig. 5A**). Expression of GATA3, a master regulator of Th2 differentiation was reduced in all boosted groups mice compared to BCG alone. This effect was enhanced when CL429 was included and delivered via the i.t. route (**Fig. 5B**). Importantly, i.t. boosting and inclusion of CL429 significantly improved levels of RORγT, a transcriptional regulator of Th17 differentiation^35^ (**Fig. 5C**). Intravenous boosting did not improve RORγT expression with or without the addition of CL429. These findings are exciting because studies in NHP models have shown that RORγT^+^ CD4^+^ T cells were associated with lower bacterial burdens in granulomas, indicating that they perform critical functions to control bacterial growth^36^. T-bet is the major transcription factor responsible for differentiating CD4^+^ T cells into a Th1 phenotype and no differences were detected in all groups compared to untreated controls (**Fig. 5D**). Transcription factor expression profiles were also consistent when total the CD4^+^ T cell population was analyzed (**Fig. S4**). Comparing the total numbers of T_EM_/T_EF_ cells expressing transcription factors, we observed higher numbers of cells in groups boosted through the i.t. route whereas i.v. boosting did not change compared with BCG only vaccinated mice (**Fig. S5**). Taken together, this data suggests that i.t. boosting with KFE8-Ag85B PNFs and CL429 combinations directs a shift in the Th profile toward a Th17 bias and i.v. boosting, in contrast, promotes a more similar distribution of all four transcription factors.

**Figure 5.**
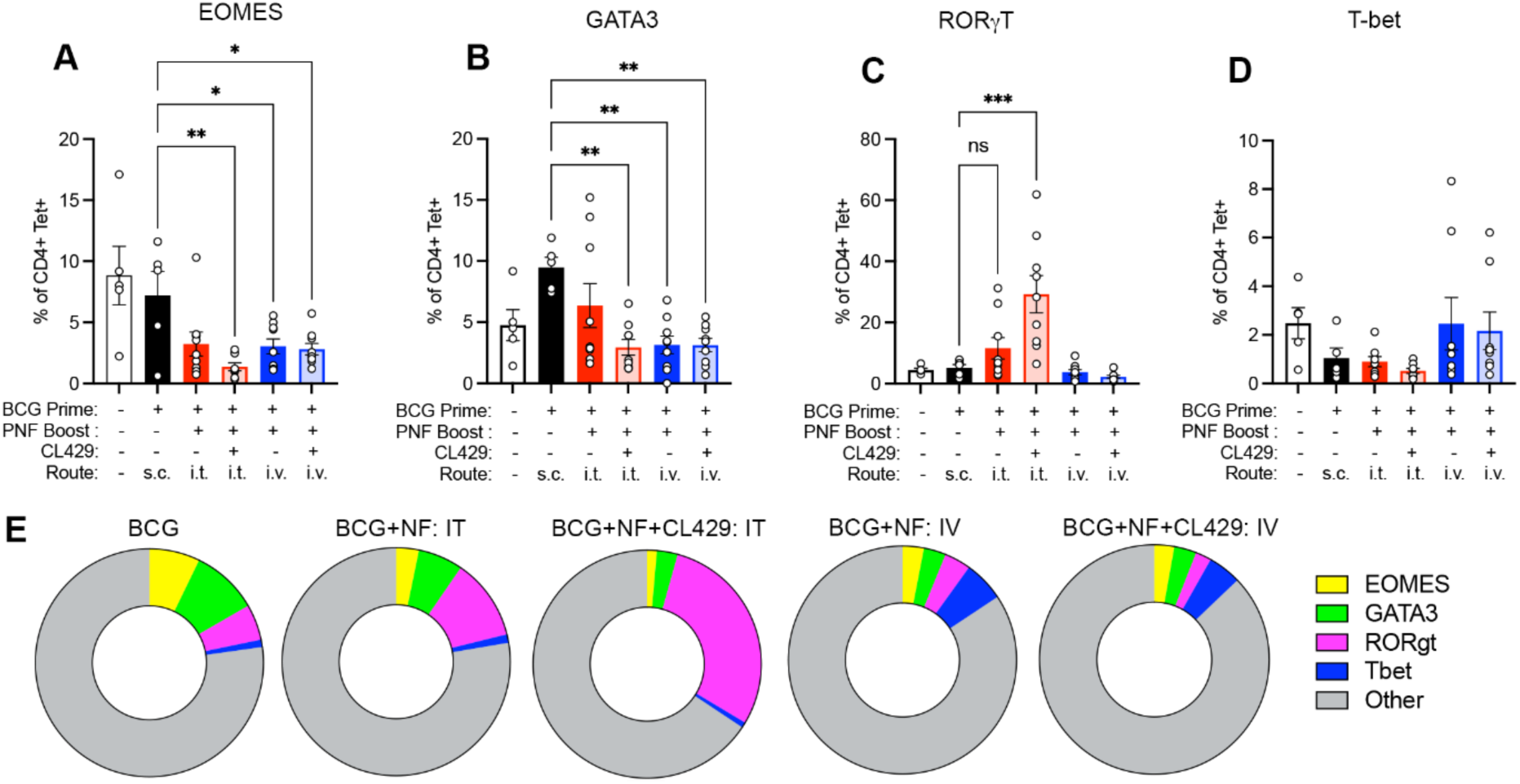
Boosting with Ag85B nanofiber/CL429 combinations drives route-dependent transcription factor bias in CD4^+^ T cells. Ag85B-specific CD4^+^ T cells were analyzed for transcription factor expression, which is represented as a percentage of the total tetramer^+^ cells. Cells were selected and gated for expression of transcription factors EOMES (**A**), GATA3 (**B**), RORgT (**C**), and T-bet (**D**). Transcription factor expression is also represented in pie charts to visualize the relative representation in each treatment group (**E**). Significant differences were determined by one-way ANOVA followed by a Benjamini Kreiger Yuketieli test to control for multiple comparisons (n = 5 in untreated and BCG only groups, n = 9 in boosted groups). *p<0.05, **p<0.005, *p<0.0005.

### CD4^+^ T cells exhibit enhanced effector cytokine profiles following PNF and CL429 boosting

While transcription factor expression is not evidence of function, it does indicate the potential of CD4^+^ T cells to express important effector molecules. To demonstrate the functional potential of memory CD4^+^ T cells in response to boosting, we analyzed the cytokine profiles in supernatants following *ex vivo* activation. IFN-γ was significantly enhanced in boosted groups but the inclusion of CL429 reduced IFN-γ production in a route-dependent fashion (**Fig. 6A**). IL-2 levels were also significantly higher in the i.v. boosted group but not influenced by CL429 (**Fig. 6B**). Consistent with RORγT expression profiles, i.t. boosting with Ag85B-KFE8 and CL429 combinations led to markedly increased IL-17A, compared to BCG alone or boosting with nanofibers alone (**Fig 6C**). A similar pattern of IL-6 production, a cytokine important in Th17 differentiation, was noted (**Fig. 6D**). Importantly, boosting via the i.v. route did not result in IL-17 or IL-6 activation. TNF levels were elevated in all boosted groups compared to mice receiving BCG alone and inclusion of CL429 enhanced TNF-α production when boosted via the i.v. route but not the i.t. route (**Fig. 6E**). Th2 cytokines IL-4, IL-5, and IL-13 were produced in abundance by CD4^+^ T cells in mice boosted nanofibers alone via the i.t. route (**Fig. 6, F to H**). Interestingly, inclusion of CL429 in mice boosted via the i.v. route led to an increase in Th2-promoting cytokines IL-4 and IL-13.

**Figure 6.**
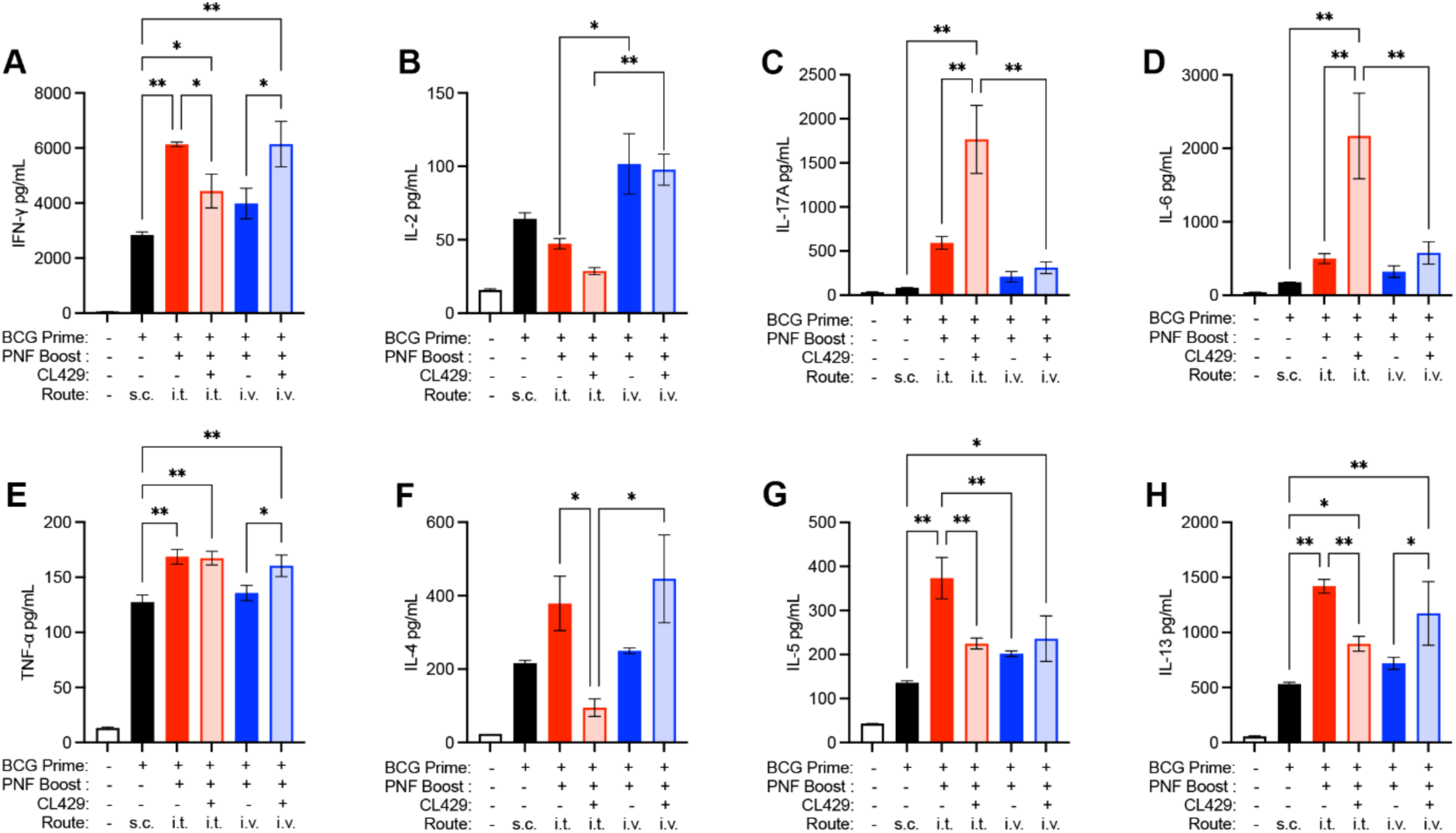
CD4^+^ T cells from lungs of boosted mice exhibit enhanced effector cytokine profiles. CD4^+^ lymphocytes were purified from disrupted lung tissue from vaccinated mice and incubated with anti-CD3/anti-CD28 antibodies for 48 hours and analyzed for cytokine content via multiplex ELISA. Analytes included IFN-γ **(A)**, IL-2 **(B)**, IL-17A **(C)**, IL-6 **(D)**, TNF-α **(E)**, IL-4 **(F)**, IL-5 **(G)**, and IL-13 **(H)**. This panel represents Th1, Th2, Th17 and inflammatory cytokines. Significant differences between treatment groups were determined by one-way ANOVA followed by a Benjamini Kreiger Yuketieli test to control for multiple comparisons (n = 2 replicates of pooled samples from 3 mouse lungs; representative of 6 samples in total). *p<0.05, **p<0.005.

Polyfunctional CD4^+^ T cells (co-secretion of 2+ cytokines per cell) are an important functional attribute of a quality immune response against *Mtb*^37^. Assessment of polyfunctional responses following boosting of BCG-primed animals with PAMP-DAMP combinations provides a potentially valuable translational tool for screening efficacy in preclinical models. To complement our cytokine from bulk supernatant, we utilized 32-plex single-cell functional proteomics platform to profile the polyfunctionality of lung CD4^+^ T cells from vaccinated mice. Purified CD4^+^ T cells from lungs of vaccinated mice were restimulated *ex vivo* and loaded into single-cell barcoded chips and cytokine production were analyzed. Data showed that untreated groups produced negligible amounts of cytokine after anti-CD3/anti-CD28 stimulation and polyfunctional T cells were detected in all boosted groups (**Fig. S6**). CD4^+^ T cells from lungs of mice primed with BCG alone and those boosted with Ag85B PNFs and CL429 via the i.t. route exhibited the highest polyfunctionality (5+ cytokines) (**Fig S6**). The functionality in the BCG group can be attributed to restimulation of a very small percentage of lung-resident cells generated from the prime. Mice receiving combination adjuvant boosters via the i.v. route also exhibited cell populations that produced 3+ or 4+ cytokines. Importantly, boosting with Ag85B nanofibers alone by either route did not promote polyfunctional populations (2 cytokines).

To assess peripheral responses, splenocytes from vaccinated mice were incubated with Ag85B-pulsed DCs for 72h and the effector cytokine profile was analyzed. BCG only and prime-boosted groups demonstrated expanded CD4^+^ T cell populations compared with the untreated controls. We performed COMPASS analysis to identify polyfunctional CD4^+^ T cells that are most likely to change in response to antigen recall. Scores derived from the analysis showed that groups boosted with nanofiber and CL429 combinations exhibited increased polyfunctionality compared to the group that received BCG only (**Fig. S7**).

### Lung-resident APCs internalize PNFs and differ in efficiency and kinetics of uptake

To determine that lung resident APC populations that take up PNFs and contribute to innate immune responses, Cy5-labeled KFE8-Ag85B nanofibers were administered via the i.t. route. Lungs were collected at 4h and 24h post inoculation to assess time-dependent effects on APCs. We analyzed a broad range of cell populations using a gating strategy^38^ that determined cell populations responsible for antigen presentation to CD4^+^ T cells (**Fig. S8**). At 4h post-delivery, alveolar macrophages (AMs, SiglecF^+^ CD11c^+^ CD11b^-^ CD64^+^) constituted approximately 35% of the Cy5^+^ cells (**Fig. 7A**), which was less than 20% at 24h. Monocytes (F4/80^+^MHCII^lo^CD11b^+^SiglecF^-^ Ly6G^-^), interstitial macrophages (IMs, CD64^+^F4/80^+^MHCII^med/lo^CD11b^+^SiglecF^-^Ly6G^-^), and DCs (MHCII^hi^ CD11c^+^ CD11b^-^ SiglecF^-^ Ly6G^-^) displayed a higher signal at 24h compared to 4h (**Fig. 7A).** Interestingly, the proportion Cy5^+^ neutrophils (CD11b^+^Ly6G^+^) also increased after 24h, while frequency of eosinophils (SiglecF^+^CD11b^+^CD11c^-^Ly6G^-^) and other myeloid cell populations (F4/80^+^CD64^+^CD11b^-^CD11c^+/-^) was reduced (**Fig. 7A**). To determine the internalization capacity, we measured the frequency of Cy5^+^ cells in each identified cell population. Nearly all AMs (∼100%) were Cy5^+^, followed by 80-90% of eosinophils and other myeloid cells and this effect was sustained 24h post-delivery (**Fig. 7B**). At 4h, ∼50% of IMs were Cy5^+^ which increased to ∼80% after 24 hours. Monocytes, DCs, and neutrophils also demonstrated enhanced uptake with time (**Fig. 7B**). To determine if the relative amount of nanofiber taken up differed between the cell subsets, we compared the MFI of Cy5^+^ between groups as a surrogate for the relative number of KFE8-Ag85B molecules. We observed that AMs exhibited the highest MFI for Cy5 at 4h and 24h, followed closely by other myeloid cells but significant differences were detected between time points (**Fig. S9**). We also analyzed whether Ag85B nanofibers drive the recruitment of innate immune cells to the lung. No significant change in any cell population was observed but differences in viability was detected between 4h and 24h. Neutrophil viability increased from 50% to 80% whereas AM viability decreased from 60% to 50% (**Fig. S10**). Eosinophils and other myeloid cells exhibited the lowest viability, which continued to decrease over time.

**Figure 7.**
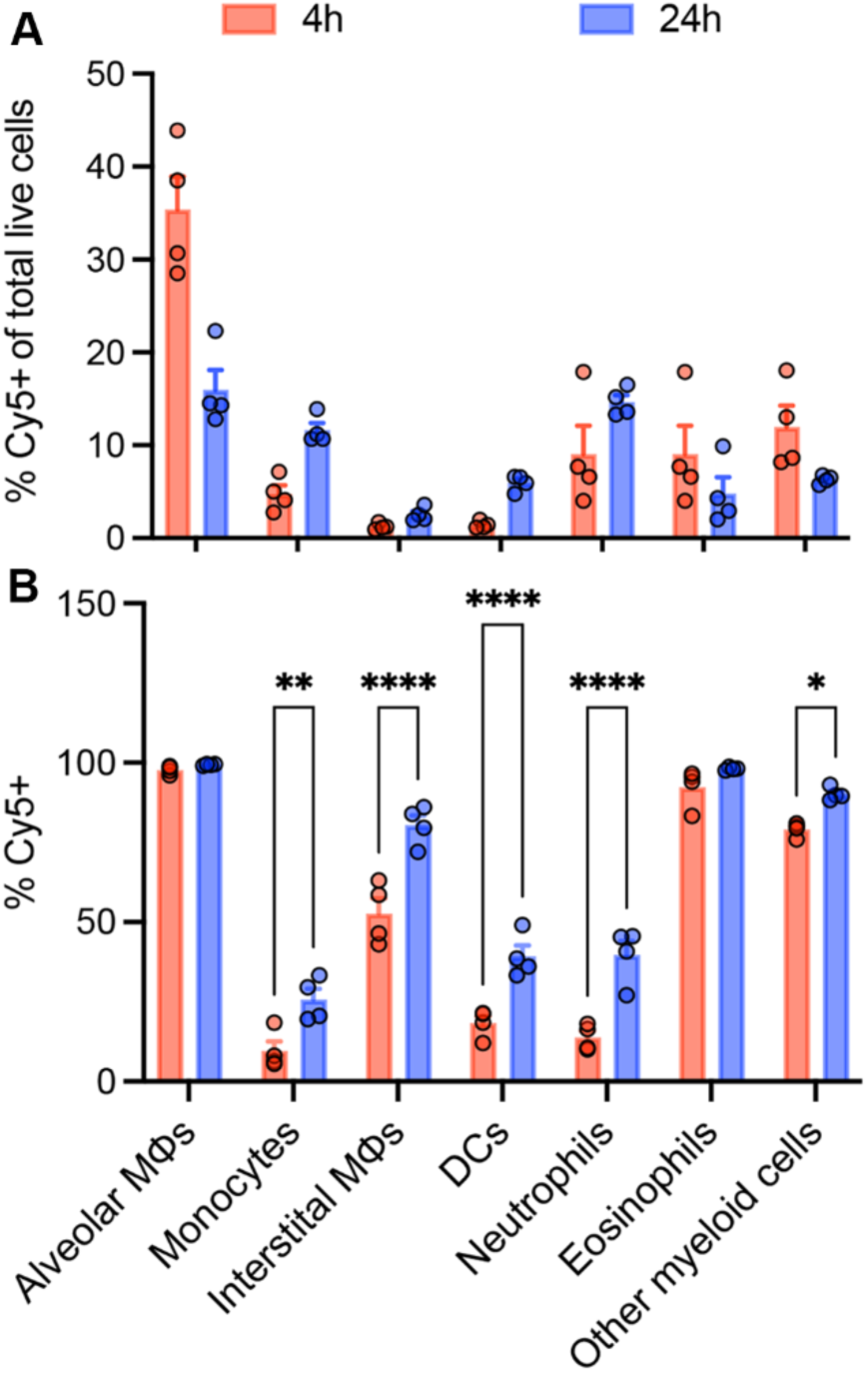
Ag85B-KFE8 nanofibers are efficiently taken up by lung APCs after i.t. delivery. Cy5-labeled peptide nanofibers were delivered via the pulmonary route and at indicated time points, Cy5^+^ cells were selected and identified through expression of surface markers described in Fig. S8. The percentage of identified APCs positive for Cy5 fluoresence are shown at 4h and 24h after treatment **(A)**. Each APC population was analyzed for the proportion of cells in that category that were Cy5^+^ **(B)**. Significant differences over time were determined by multiple t tests followed by a correction for multiple comparisons using the Holm-Sidak method (n = 4). *p<0.05, *p<0.005, ****p<0.0005.

### Lung APCs exhibit increased co-stimulatory capacity following PNF uptake

To assess immune cell activation, we selected for three important professional APC populations including interstitial macrophages (IM), alveolar macrophages (AM), and dendritic cells (DC) using CD11c, CD11b, and MMR (CD206) markers (**Fig. 8A**). Four hours after i.t. delivery of Cy5-KFE8-Ag85B, ∼2.5% of the total viable cell population in the lungs were Cy5^+^ which increased to 4% after 24h (**Fig. 8B**). We compared the relative expression of CD80 and CD86 on untreated APCs, Cy5^+^ APCs after 4h and 24h of exposure. CD80 was greatly increased in IMs and DCs at both time points compared with the untreated controls (**Fig. 8C**). AMs and DCs exhibited a significant upregulation of CD80 at 4h, but with less intensity (**Fig. 8D and 8E**). However, there was no significant difference in expression between 4h and 24h. CD86 expression was also significantly increased in IMs and DCs 4 hours after treatment (**Fig. 8F and 8H**). CD86 expression by AMs did not increase compared with untreated controls and was significantly reduced after 24h (**Fig. 8G**). Taken together, these findings suggest that vaccination with Ag85B nanofibers results in recruitment of innate immune cells to the lungs and induces activation of professional APCs.

**Figure 8.**
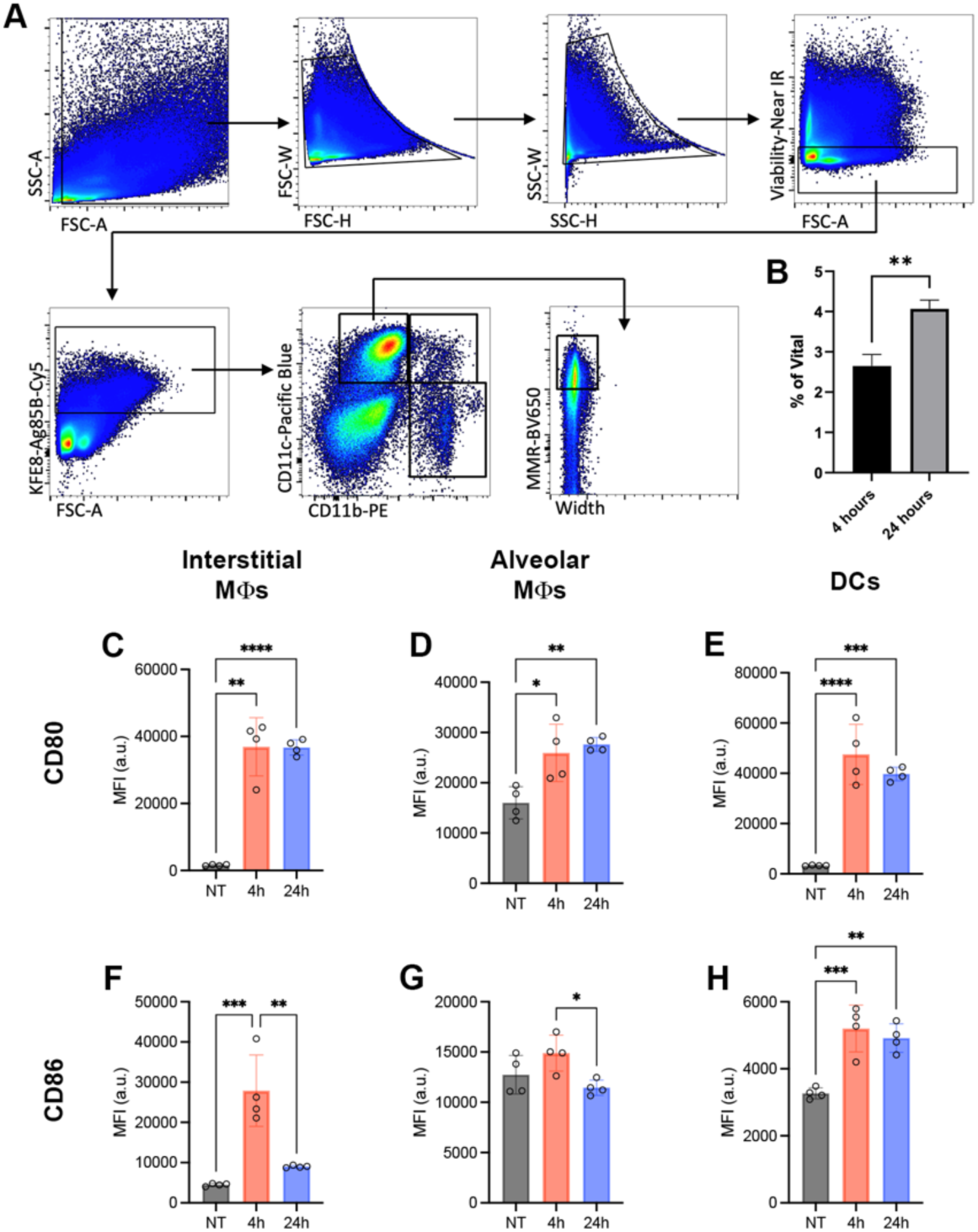
APCs exhibit increased expression of co-stimulatory molecules following uptake of KFE8-Ag85B nanofibers. Lungs were collected from mice 4h and 24h after pulmonary administration of Cy5^+^ labeled KFE8-Ag85B nanofibers. The lung tissue was processed for flow cytometric analysis Following a hierarchal gating strategy, single cells were selected through doublet exclusion, followed by the selection of viable cells based on exclusion of cells that positively stained for viability dye. Then Cy5^+^ were selected as NF^+^ cells. The expression of CD11b, CD11c, and MMR (CD206) was used to putatively identify interstitial macrophages (CD11b^+^CD11c^+^), alveolar macrophages (CD11c^+^CD11b^-^MMR^+^), and DCs (CD11c^+^CD11b^-^ MMR^-^) (**A**). The frequency of NF^+^ cells is represented as a percentage of the total viable cell population in disrupted lung tissue 4h and 24h after i.t. delivery (**B**). Interstitial macrophages, alveolar macrophages, and DCs were analyzed for their expression of CD80 (**C-E**) and CD86 (**F-H**), which is represented as the MFI in untreated animals and 4 and 24 hours after treatment. Significant differences between treatment groups were determined by one-way ANOVA followed by a Tukey test to control for multiple comparisons (n = 4). *p<0.05.

## DISCUSSION

In the last decade, T_RM_ cells have emerged as an important subset of non-circulating memory T cells that provide potent surveillance at the mucosal barrier. T_RM_ cells remain localized in their home tissues due to a combination of adhesion molecules and homing mechanisms that favor trafficking toward sites of inflammation and retention in tissues, and resist egress to lymphoid tissues. Lung T_RM_ and T_EM_ are associated with lower infection risk and enhanced control of active TB. Adoptive transfer of total lung T cells following mucosal vaccination with *Mtb* antigens protects against TB^13^ and mucosal vaccination with attenuated strains of *Mtb* is highly effective in NHPs^14^. Other strategies to induce lung T_RM_ cells have focused on boosting BCG vaccinated mice with subunit vaccines. *Bacillus subtilis* spores, Sendai virus vector, cationic liposomes, and nanoparticles bearing *Mtb* antigens have been shown to be effective at boosting BCG and enhancing protection^15^. These studies provide experimental evidence that circulating *Mtb*-specific T cells are associated with protection and vaccines that can boost lung T_RM_ cell populations are effective at controlling *Mtb* infection.

Here, we demonstrate that the route of booster delivery and inclusion of PAMP agonists can affect the memory precursor populations, the effector cytokine profiles, and the transcription factor bias that determines the differentiation of antigen-specific CD4^+^ T cells. Boosting BCG-primed mice using DAMP-inducing PNFs via the i.t. route expands CD4^+^ and CD8^+^ memory precursor cell populations which were further enhanced when CL429, a PAMP agonist, was included. Also, i.t. boosting generated substantial numbers of antigen-specific T cells and these cells were almost exclusively KLRG1^-^ CX3CR1^-^ memory precursor cells. Parenchymal lung CD4^+^ T cells in the setting of *Mtb* infection are predominantly KLRG1^-^ CX3CR1^-^ and produce less IFN-γ than circulating memory populations^8^, but are highly proliferative and survive longer than their intravascular KLRG1^+^ counterparts^24^. KLRG1^-^ CX3CR1^-^ cells have also been shown to express a high frequency of CXCR3, an important homing marker and commonly found on T_RM_ cells^20,39^. Interestingly, CXCR3 expression was greatest in the i.t. vaccinated mice, suggesting homing of antigen specific CD4^+^ T cells to the lung parenchyma.

Prior studies have shown that KFE8 nanofibers are a strong inducer of Th2 cytokines (IL-4, IL-5) responsible for driving humoral responses but can induce a mixed Th1/Th2 response when combined with TLR agonists. One study using Q11-based peptide nanofibers showed uptake and presentation of MHC-II peptide antigens on lung CD103^+^ and CD11b^+^ DCs, up-regulation of CD80, and a predominantly T_H_17 response, with minimal T_H_1 or T_H_2 responses in the absence of exogenous adjuvants^40^. In this study, the addition of CL429 to Ag85B-KFE8 nanofibers led to reduction of important Th1 (IFN-γ and IL-2) and Th2 (IL-4, IL-5, and IL-13) cytokines, reflecting a state of heightened inflammation, as demonstrated by the increase in IL-6. In contrast, i.v. boosting in combination with CL429 enhanced Th1 and Th2 cytokines, while IL-17 and IL-6 remained relatively low. These results indicate that when delivered via the i.t. route, KFE8 nanofibers drive Th1 and Th2 responses, but require the addition of CL429 to generate IL-17 responses. Intravenous delivery of Ag85B nanofibers resulted in only a moderate accumulation of T lymphocytes in the lung. Although the lung has a dense capillary bed, it is possible that the nanofibers do not extravasate easily to enable capture by pulmonary APCs and are taken up for processing at other tissue sites. Future studies characterizing the biodistribution of peptide nanofibers delivered via the i.t. versus i.v. route will provide better insights into their in vivo fate and subsequent evolution of memory T cell responses. Collectively, we demonstrate that i.v. boosting with peptide nanofiber vaccines can generate lung-resident immune responses, as well as a shift in Th bias that differs from mucosal delivery of peptide nanofibers.

These results are supported by analyses of cytokine secretion in lung supernatants where IL-17A and IL-6 production was highest in mice that received i.t. boosts and drastically enhanced by the addition of CL429. Analysis of transcription factor expression demonstrated a decrease in EOMES and GATA3, the transcription factor driving Th2 responses. Consistent with previous reports on mucosal vaccination^40,41^, RORγT was upregulated in groups that received pulmonary boosts, especially in Ag85B-specific CD4^+^ T cells, indicating a shift toward Th17 bias. This is supported by the concurrent reduction in EOMES expression, known to be increased in inflamed tissues^34,42^ and suppress RORγT and IL-17 production^34,43^. In contrast, EOMES has been shown to promote CD8^+^ T cell exhaustion^44^ and CD4^+^ IFN-γ production in the absence of T-bet^42,43^. Surprisingly, i.v. vaccination did not induce pronounced changes in transcription factor expression that we observed with i.t. vaccination, although T-bet was moderately upregulated, suggesting a slight bias toward Th1 type.

*Mtb* subverts the functions of lung APCs, thereby limiting efficient T cell priming and allowing an infection to be established^45,46^. Consistent with this observation, *Mycobacteria spp* including BCG have been shown to downregulate the expression of CD80 and CD86 on APCs^47,48^. Closing the communication gap between APCs and T cells is imperative in establishing immune memory. Our data show that APCs that internalize KFE8 nanofibers express co-stimulatory molecules CD80 and CD86 necessary for efficient T cell priming. This is consistent with other studies demonstrating that intraperitoneal vaccination with peptide nanofibers upregulates the expression of CD80 and CD86 and highlights the adjuvanting potential of peptide nanofibers independent of the route of administration^49^. CD80 and CD86 expression on AMs and DCs differentially and synergistically regulate T cell responses in the lung, through PI3 kinase and other signaling pathways that activate proliferation, augment survival, and promote cytokine secretion^50,51^. A study utilizing *Mtb* secretory antigen-primed DCs demonstrated that stimulation with anti-CD80 monoclonal antibodies induced IFN-γ production and upregulated CD86 expression, which in turn modulated antigen-specific T cell recall responses through upregulation of IL-10^52^. In contrast, both CD80 and CD86 have also been shown to promote Th2 cytokine production in T cells^53^. Our results identify the lung APCs that are targeted by PNF to promote the generation of highly activated populations of T cells in the lungs of boosted mice.

We observed differential uptake of nanofibers by lung APCs where AMs exhibited the highest internalization not only in terms of numbers but also relative quantity of nanofibers. The reduction in numbers and viability of Cy5^+^ AMs after 24h presumably indicates trafficking to draining lymph nodes or cell death leading to DAMP release^33^. DCs and neutrophils exhibited a time-dependent increase in recruited numbers and internalization, likely from phagocytosing smaller aggregates of nanofibers or nanofibers associated with dead cell debris. DCs are powerful potentiators of T cell activation and adoptive transfer of antigen-loaded DCs overcomes antigen presentation bottlenecks in TB^46,54^. Additionally, DCs can present antigen from dead cells, promoting cross-presentation^55,56^. Sterile inflammation increases neutrophil chemotaxis, which may also internalize peptide nanofibers^57^. The involvement of eosinophils is interesting and they respond to DAMPs, such as IL-33, and promote a Th2 bias^58^ as well as activation of other innate immune cells^59^. Additionally, it was recently reported that eosinophils may play a protective role in the immune response to pulmonary TB, in part due to degranulation activity in *Mtb* lesions and granulomas^60^. The category of other myeloid cells was comprised of cell populations that could not be appropriately defined were ruled out of other populations based on their lack of consistent expression of markers other than F4/80 and internalization of Cy5-labeled nanofibers.

## CONCLUSION

In conclusion, we demonstrate that combination adjuvants composed of Ag85B-KFE8 nanofibers and agonist of the TLR2/NOD2 signaling cascade (CL429) are a robust platform for boosting BCG-primed immunity and generating lung T_RM_ cells with robust effector functions. Following pulmonary administration, lung APCs efficiently internalize peptide nanofibers and upregulate important co-stimulatory markers that drive T cell priming and activation, as we have previously described for extrapulmonary APC^31^. The route of immunization with Ag85B-KFE8+CL429 differentially activates key transcription factors that determine functional specialization of Th cells, resulting in a Th17 bias when delivered via a mucosal route or a Th1 bias when delivered intravenously.

These studies in a pre-challenge model pave the way for future work to establish protective efficacy of in mouse models of *Mtb* challenge and generate multicomponent vaccines with controlled stoichiometric ratios of antigens, nanofibers, and adjuvants. Application of multifactorial experimental approaches such as Design of Experiments (DOE) and Response Surface Methodology (RSM) to tune vaccine composition will enable use of a reductionist approach to determine how each PAMP agonist affects DC activation, antigen-presentation, and T cell function individually and in combination with peptide nanofibers. Our findings demonstrate the potential for heterologous booster vaccines composed of peptide nanofibers and PAMP agonist combination adjuvants to engage innate and adaptive immunity for generating T_RM_ cells with established roles in protection against TB and other respiratory diseases.

## Notes

### Competing Interest Statement

The authors have declared no competing interest.

