## Supplementary Information for "DAMP-inducing Peptide Nanofibers and PAMP Combination Adjuvants Boost Functional Lung Tissue-resident Memory CD4^+^ T Cell Responses"

**Figure S1**. Gating strategy for effector, memory, and antigen specific T cells	S2

**Figure S2**. Frequency of KLRG^-^CX3CR1^-^CD4^+^T_EF_ cells	S3

**Figure S3**. Activation of CD8^+^T cells	S4

**Figure S4**. Transcriptomic profile of total CD4^+^T cells	S5

**Figure S5**. Total numbers of T_EM_/T_EF_ cells expressing transcription factors	S6

**Figure S6**. Single cell secretome profile of lung CD4^+^T cells	S7

**Figure S7**. Peripheral CD4^+^T cell responses and cytokine production	S8

**Figure S8**. Gating strategy for lung-resident APCs	S9

**Figure S9**. Time-dependent uptake of nanofibers by immune cells	S10

**Figure S10**. Viability of lung APCs	S11

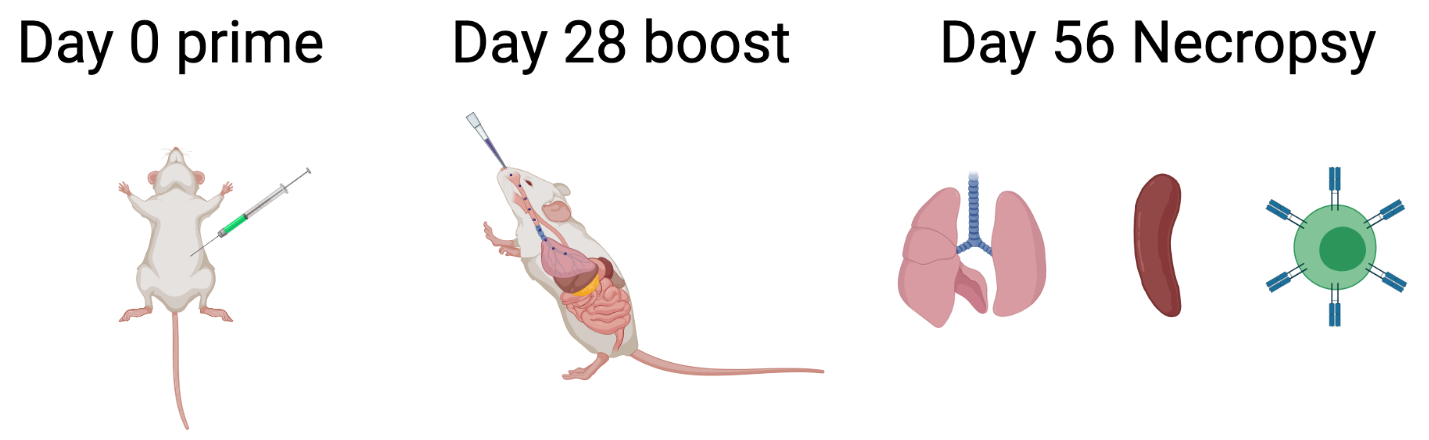

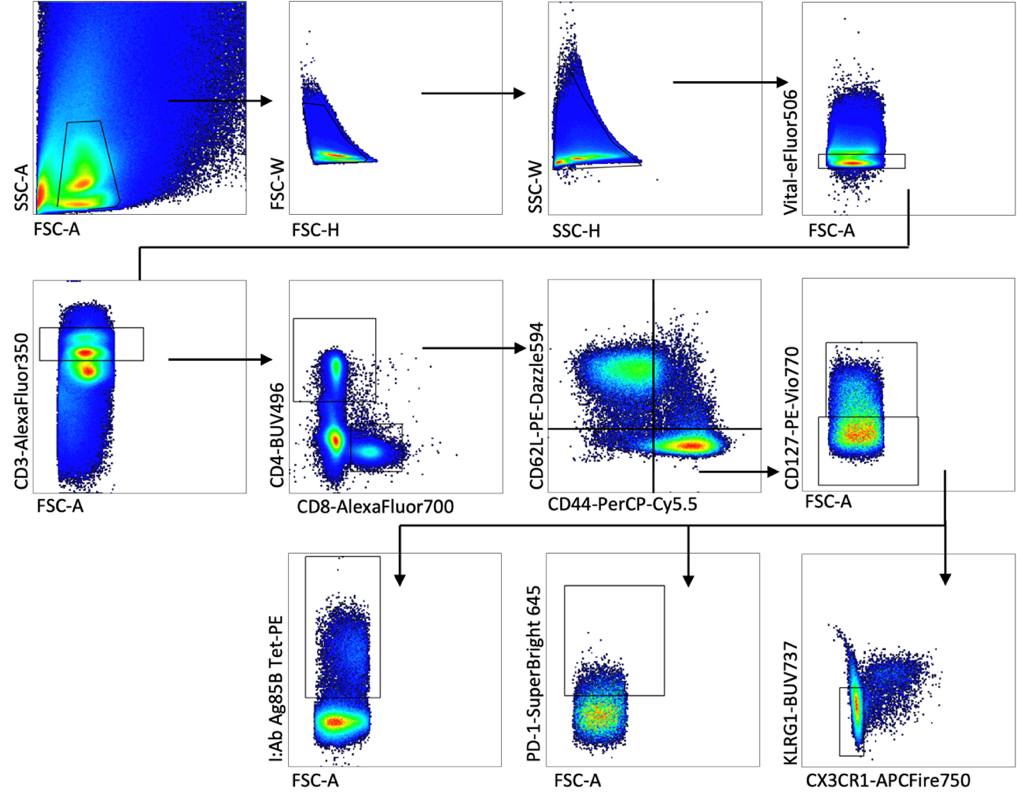

**Figure S1. Gating strategy for effector, memory, and antigen-specific T cells.** BCG-primed mice were given two boosts four and seven weeks post prime. Boosts with and without adjuvant were delivered either delivered via i.t. or i.v. route. Mice were euthanized 7, 8, and 9 days after the second boost (3 mice each day). Lungs and spleen were collected from each animal and assayed using flow cytometry and single cell secretome analysis. Spleens were used in antigen recall assays and cytokine content was analyzed using multiplex ELISA. Cells from lung homogenate were gated on CD4^+^CD3^+^ lymphocytes and CD8^+^CD3^+^ lymphocytes (not shown). CD44 and CD62L were used to identify subsets of activated T cells, known here as central memory (T_EM_) and effector memory (T_EM_) or effector T cells (T_EF_), which are characterized as CD44^hi^CD62L^hi^ and CD44^hi^CD62L^lo^ respectively. The T_EM_/T_EF_ subset was further evaluated for expression of CD127 (IL-7Ra), which differentiates T_EM_ (CD127^+^) from T_EF_ (CD127^-^). Terminally differentiated T cells are distinguished by the elevated expression of KLRG1 and CX3CR1, while long-lived memory T cell populations lack these markers. T_EM_ and tissue-resident memory T cells (T_RM_) also express CXCR3 with greater intensity.

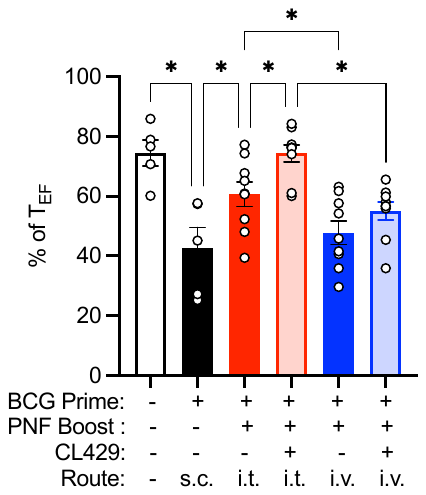

**Figure S2. Pulmonary boost with KFE8-Ag85B alone in BCG-primed mice increases the frequency of KLRG^-^CX3CR1^-^CD4^+^T_EF_ cells.** Cells that are fated to become memory T cells do not express KLRG1 or CX3CR1 and the percentage of CX3CR1^-^ KLRG1^-^ cells are expressed as a percentage of CD4^+^ T_EF_ cells. Statistical significance was determined by one-way ANOVA followed by a Benjamini Krieger Yuketieli test to control for multiple comparisons (n = 5 in untreated and BCG only groups, n = 9 in boosted groups). *p<0.05

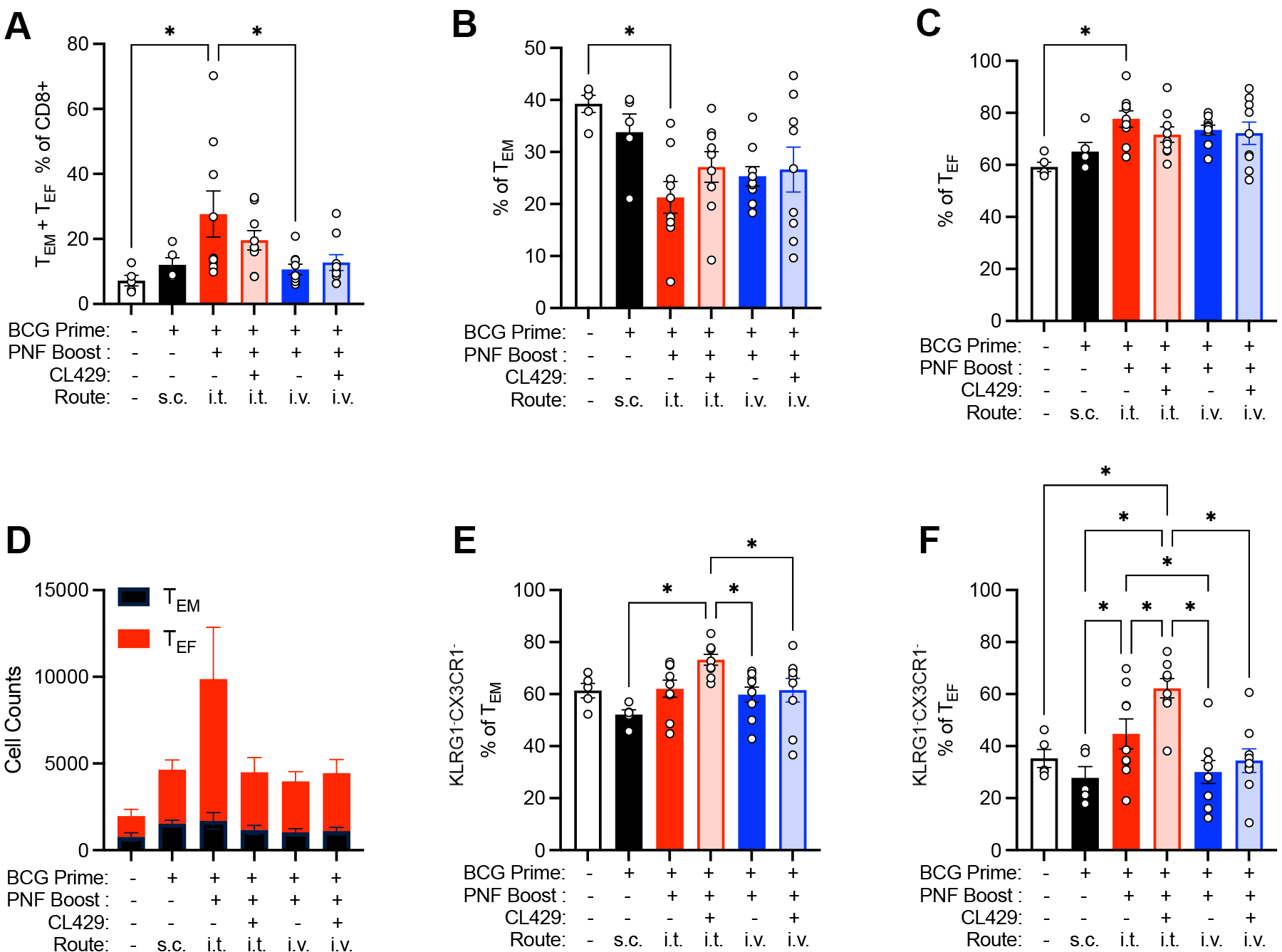

**Figure S3. Pulmonary boost with KFE8-Ag85B in BCG-primed mice increases the frequency of CD8^+^ T_EM_ and T_EF_ cells.** BCG-primed mice were given two boosts four weeks and seven weeks later with Ag85B-KFE8 nanofibers with or without CL429 adjuvant. Booster doses were delivered via the i.t. or i.v. route. Mice were euthanized after the second boost and lungs were collected for flow cytometry **(A)**. CD8^+^ T_EM_ and T_EF_ cells were identified as CD44^hi^ CD62L^lo^ and further differentiated based on their expression of CD127. T_EM_ (CD127^+^) cells (**B**) and T_EF_ (CD127^-^) (**C**) cells are shown as a percentage of the total CD4^+^ T cell population. Total counts of T_EM_ and T_EF_ cells are shown in stacked plots (**D**). CD8^+^ T_EM_ cells lacking expression of KLRG1 and CX3CR1 are represented as a percentage of total CD8^+^ T_EM_ (**E**) and T_EF_ (**F**) populations. Statistical significance was determined by one-way ANOVA followed by a Benjamini Krieger Yuketieli test to control for multiple comparisons (n = 5 in untreated and BCG only groups, n = 9 in boosted groups). *p<0.05

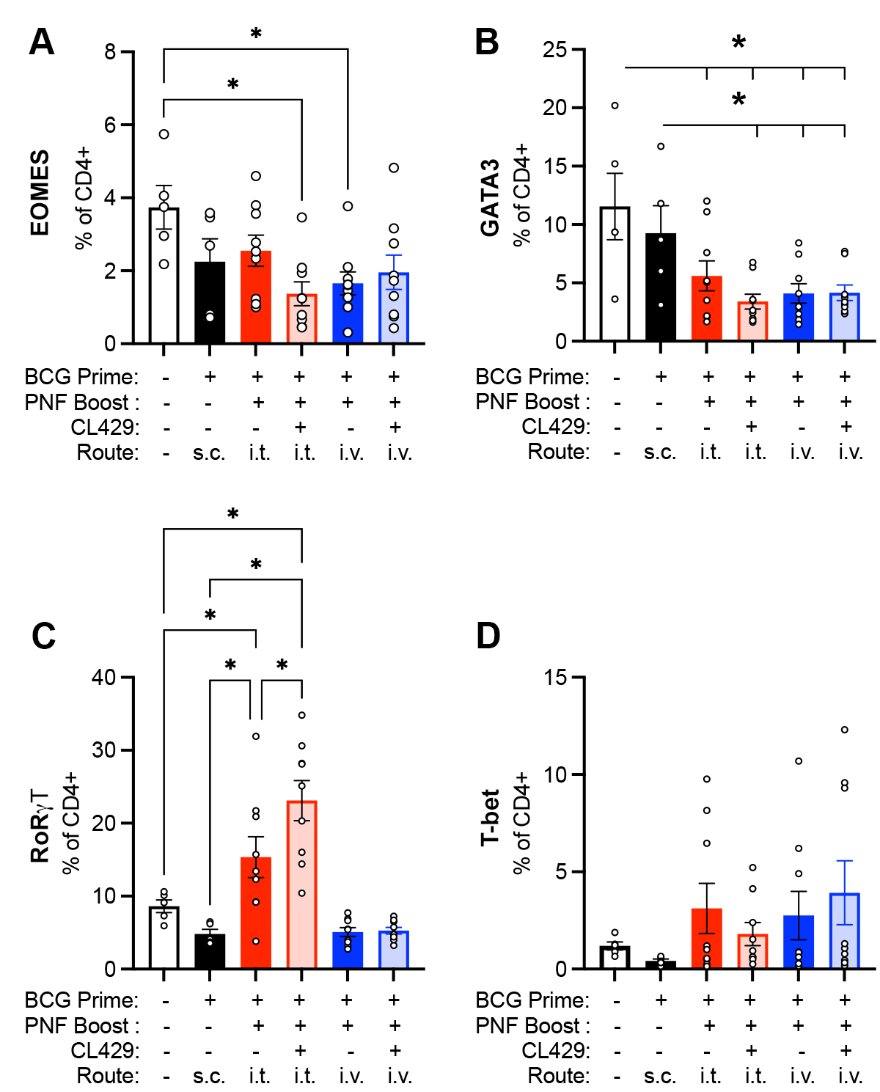

**Figure S4. Boosting with Ag85B nanofiber and CL429 combinations drives route-dependent transcription factor bias in CD4^+^ T cells.** Transcription factor-positive CD4^+^ T cells were analyzed as a percentage of the total CD^+^T cells. Cells were selected and gated for expression of transcription factors EOMES (**A**), GATA3 (**B**), RORγT (**C**), and T-bet (**D**). Significant differences were determined by one-way ANOVA followed by a Benjamini Kreiger Yuketieli test to control for multiple comparisons (n = 5 in untreated and BCG groups, n = 9 in boosted groups). *p<0.05.

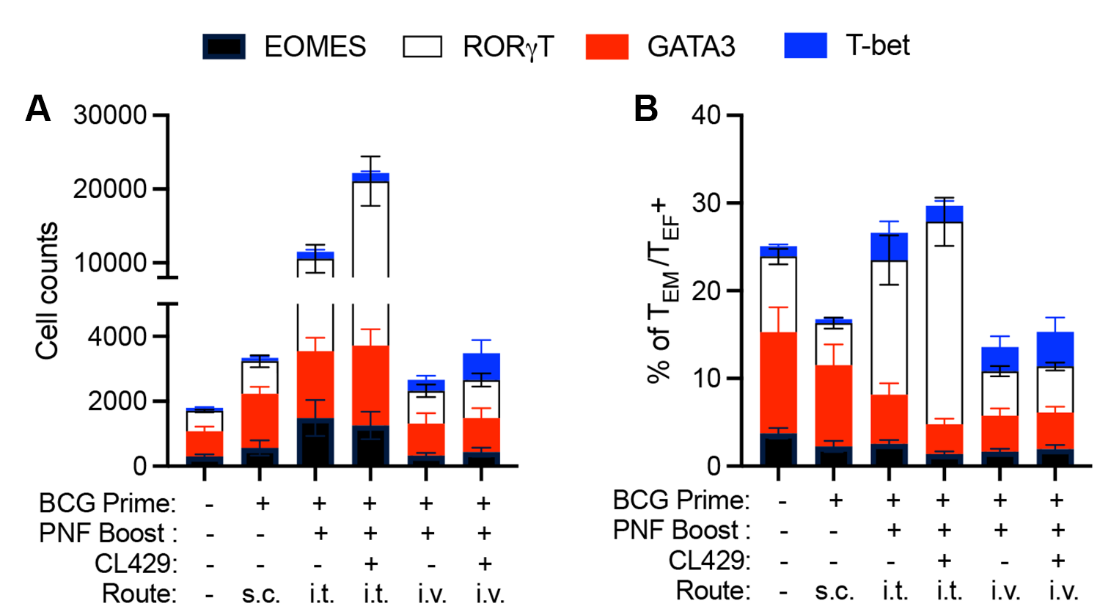

**Figure S5. Total numbers of T_EM_/T_EF_ cells expressing transcription factors.** CD4^+^ T_EM_/T_EF_ cells were selected by flow cytometry and gated for expression of transcription factors EOMES, GATA3, RORγt, and T-bet. Transcription factor expression is represented in stacked columns as total cell counts **(A)** and as a percentage of CD4^+^ T_EM_/T_EF_ cells **(B)**.

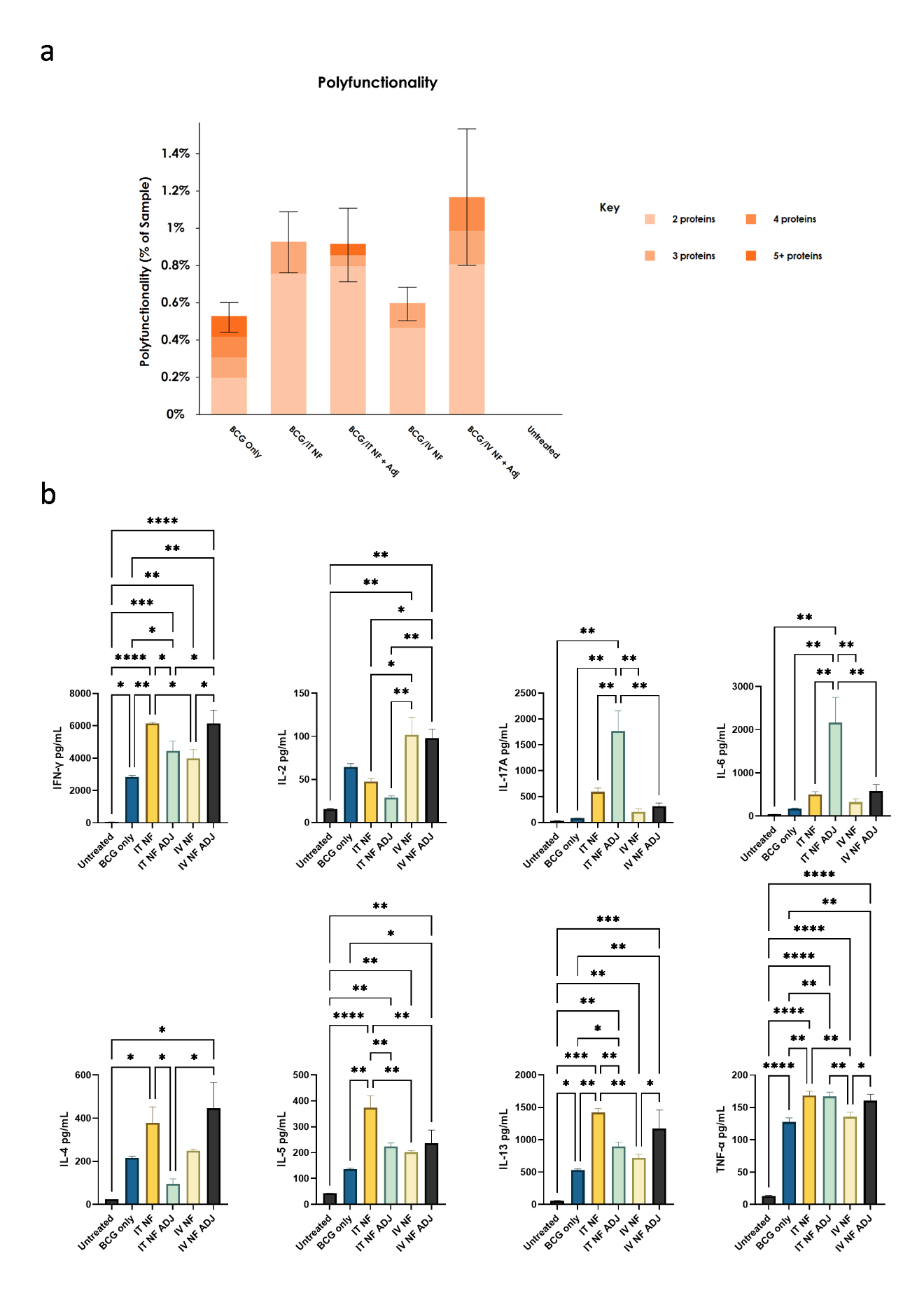

**Figure S6. CD4^+^ T cells from lungs of vaccinated mice exhibit enhanced effector cytokine profile.** CD4^+^ lymphocytes were purified from disrupted lung tissue from vaccinated mice and incubated with anti-CD3/anti-CD28 antibodies for 48 hours. The single-cell adaptive immune secretome was analyzed using the Isoplexis technology (n = 3). Cells expressing two or more cytokines were plotted as a percentage of total cells analyzed and categorized based on the number of cytokines expressed. Significance was determined using the IsoSpeak software.

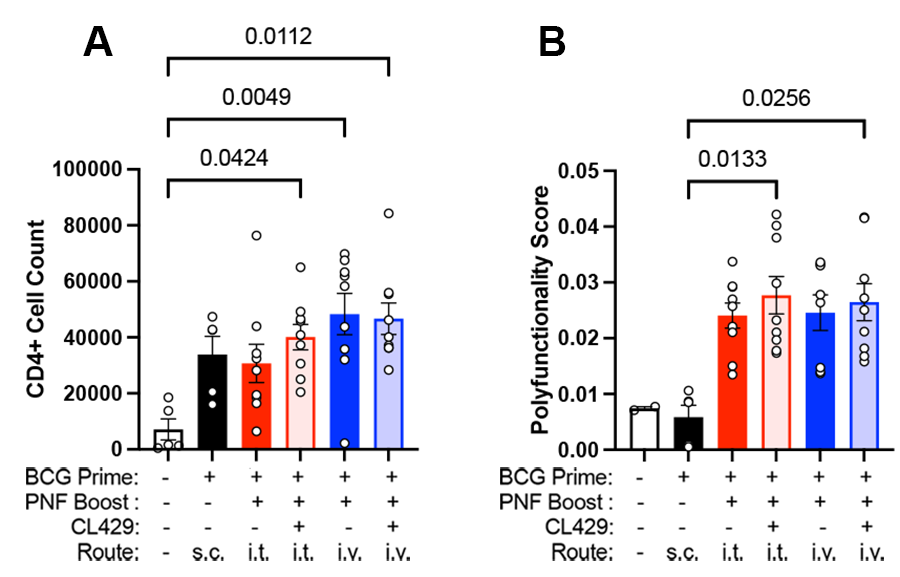

**Figure S7.** Splenic CD4^+^T cells exhibit increased polyfunctionality following i.v. delivery of Ag85B-KFE8 in BCG-primed animals. CD4^+^T cells counts are displayed for each treatment group following 72 hours of ex vivo stimulation of splenocytes in the presence of cognate antigen, Ag85B **(A)**. Cytokine expression was measured using intracellular flow cytometry and COMPASS was used to determine functional subsets of CD4^+^T cells which are likely to be increased in the stimulated sample compared with the vehicle control **(B)**. Polyfunctionality scores were derived from the COMPASS analysis and plotted for each group.

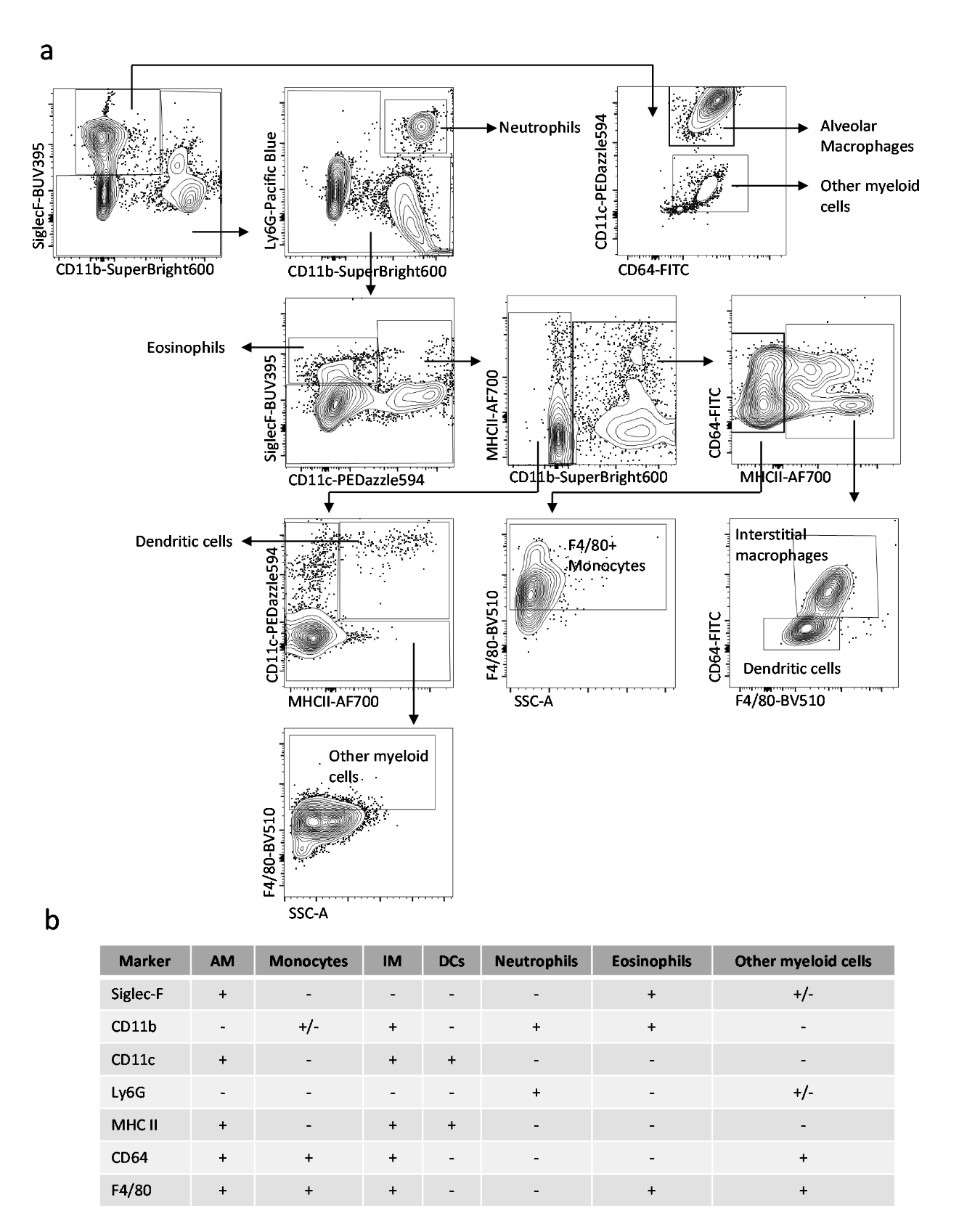

**Figure S8. Gating strategy selects for a diverse array of murine lung APCs.** Cell populations in murine lung were identified and categorized based on surface expression of markers identifying alveolar macrophages, interstitial macrophages, monocytes, DCs, neutrophils, and eosinophils. Remaining F4/80^+^ cells were categorized as other myeloid cells and all other cells that could otherwise not be identified were categorized as other **(A)**. Complete description of surface marker expression in each population **(B)**.

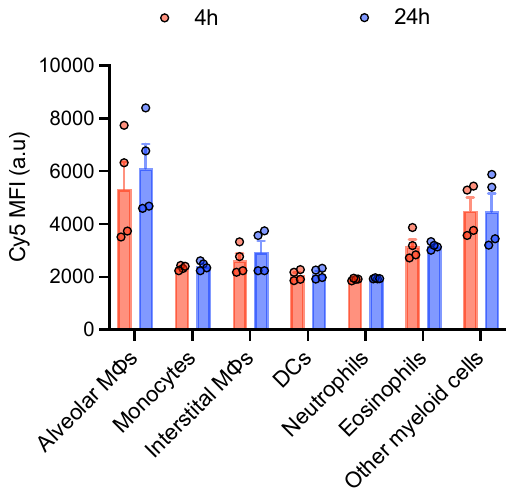

**Figure S9. Time-dependent uptake of Cy5-labeled KFE8-Ag85B by lung APCs populations.** Cy5-labeled peptide nanofibers were delivered via the pulmonary route and at indicated time points, Cy5^+^ cells were selected and identified through expression of surface markers described in Fig. 10. Each APC population was analyzed for the proportion of cells in that category that were Cy5^+^ reflecting the relative amount internalized by each cell population.

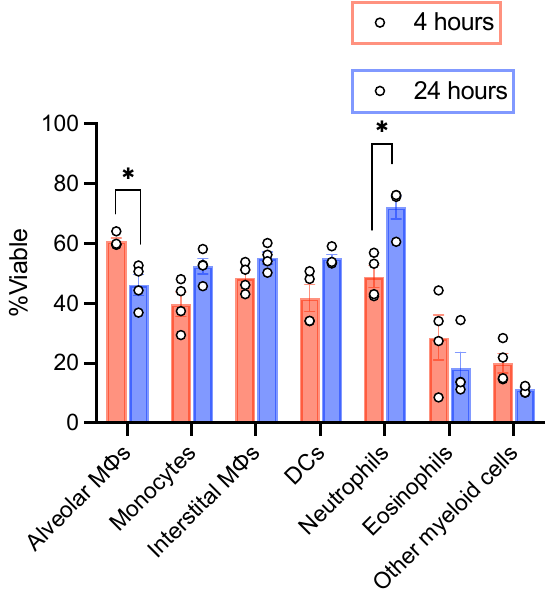

**Figure S10.** **Viability of APCs populations following Cy5-labeled KFE8-Ag85B uptake.** Cy5-labeled peptide nanofibers were delivered via the pulmonary route and at indicated time points, Cy5^+^ cells were selected and identified through expression of surface markers described in Fig. 10. The viability of each APC population was determined at 4 and 24 hours after immunization. Significant differences over time were determined by multiple t tests followed by a correction for multiple comparisons using the Holm-Sidak method (n = 4). *p<0.05.
